# Rapid, automated and experimenter-free assessment of cognitive flexibility reveals learning impairments following recovery from activity-based anorexia in female rats

**DOI:** 10.1101/2022.11.15.516539

**Authors:** Kaixin Huang, Laura K Milton, Harry Dempsey, Stephen J Power, Kyna-Anne Conn, Zane B Andrews, Claire J Foldi

## Abstract

Anorexia nervosa (AN) has among the highest mortality rates of any psychiatric disorder and is characterised by cognitive inflexibility that persists after weight recovery and contributes to the low rates of recovery. What remains unknown is whether cognitive inflexibility predisposes individuals to AN, a question that is difficult to determine from human studies. Our previous work using the most well-established animal model of AN, known as activity-based anorexia (ABA) identified a neurobiological link between cognitive inflexibility and susceptibility to pathological weight loss in female rats. However, testing flexible learning prior to exposure to ABA in the same animals has been thus far impossible due to the length of training required and the necessity of daily handling, which can itself influence the development of ABA.

Here we describe experiments that validate and optimise the first fully-automated and experimenter-free touchscreen cognitive testing system for rats (n=20) and use this novel system to examine the reciprocal links between reversal learning (an assay of cognitive flexibility) and weight loss in the ABA model (n=60). Firstly, we show substantially reduced testing time and increased throughput compared to conventional touchscreen testing methods because animals engage in test sessions at their own direction and can complete multiple sessions per day without experimenter involvement. We also show that, contrary to expectations, cognitive inflexibility does not predispose rats to pathological weight loss in ABA but instead that rats subsequently susceptible to weight loss performed better on the reversal learning task. Intriguingly, we show reciprocal links between ABA exposure and cognitive flexibility, with ABA exposed (but weight recovered) rats performing much worse that ABA naïve rats on the reversal learning task. On the other hand, animals that had been trained on reversal learning were better able to resist weight loss upon subsequent exposure to the ABA model. We also uncovered some stable behavioural differences between ABA susceptible versus resistant rats during touchscreen test sessions using machine learning tools that highlight possible predictors of anorectic phenotypes.

These findings shed new light on the relationship between cognitive inflexibility and pathological weight loss and provide a robust target for future studies using the ABA model to investigate potential novel pharmacotherapies for AN.

## Introduction

Cognitive flexibility refers to the capacity to modify behavioural choice to meet the demands of a changing environment and is crucial for selecting appropriate responses based on context and circumstance [1]. Impairments in cognitive flexibility are common to a range of psychiatric illnesses including schizophrenia [2, 3], obsessive compulsive disorder [4], and addictive disorders [5, 6], which are characterized by stereotypical patterns of rigid behaviours that persist despite negative consequences, ultimately impacting decision-making. Individuals with a current or previous diagnosis of anorexia nervosa (AN) also exhibit rigid behaviours, especially surrounding illness-relevant stimuli such as feeding and exercise [7–11]. While impaired cognitive flexibility is most severe in patients acutely ill with AN and likely contributes to perpetuating the condition [7, 10], the persistence of inflexible behaviour following weight recovery and in unaffected sisters of patients with AN suggests that it is involved in the aetiology of the disorder [10–12]. What remains to be determined is whether cognitive inflexibility itself predisposes individuals to develop AN and could be used as a biomarker to predict illness onset or severity in individuals at risk. Moreover, a detailed understanding of the neurobiology underlying an inflexibility that persists after weight recovery in individuals with AN is imperative to develop novel pharmacotherapies that can aid in long term recovery [13–15].

While the premise that cognitive rigidity is a fundamental trait of AN is well-accepted, measures of cognitive flexibility in patient populations are prone to inconsistent findings between studies, a complication that is likely amplified by large discrepancies in participant demographics and experimental approaches [16]. It is also difficult to determine from human studies the neurobiological mechanisms that precede the development of AN that could act as targets for early intervention. The question then arises - how can we assess the neural mechanisms of cognitive flexibility in animal models that adequately captures the clinical presentation in AN patients? Rodent models that incorporate key aetiological features, such as the most well-established animal model of AN known as activity-based anorexia (ABA), have been instrumental for identifying the specific neural circuits that contribute to disordered cognitive functioning [17]. Additionally, the last decade has witnessed an explosion in the availability of innovative tools including optogenetics [18], chemogenetics [19] and calcium imaging [20], to manipulate and record neural activity in freely behaving animals. These approaches give an unprecedented ability to answer questions about the relationship between brain function and behaviour relevant to a range of human disorders, including AN. However, the interest in new techniques to modify and record brain function has not been matched with adequate enthusiasm regarding the quantification and analysis of behavioural outputs that are critical for assessment of these relationships.

With this in mind, the study of cognition and behaviour in rodents has benefited in recent years from advances in technology that have increased the translational capacity of rodent models of human pathologies [21, 22]. A major contribution to improving translation has been the incorporation of touchscreens displaying visual stimuli in rodent test batteries that closely mimic those used for human cognitive testing [22], which improves standardization and interpretation of data. However, touchscreen testing in rodents has thus far required significant time and experimenter intervention to transfer subjects to and from the testing chamber. Indeed, it is well known that experimenter involvement influences experimental outcomes, particularly so for behavioural studies - including those involving the ABA model, in which the outcomes are known to be influenced by experimenter handling [23]. Along with stress from handling, which varies between experimenters and therefore differentially impacts upon task performance [24–26], manual transfer to test chambers at times that suit the experimenter is insensitive to the current motivational state of the animal and disrupts normal social behaviour. Thus, while the wide adoption of touchscreen cognitive testing has already yielded substantial benefits for behavioural neuroscience, the next frontier lies in the automation of the role of the experimenter in gatekeeping touchscreen access [27, 28].

One approach has been to relocate the operant testing modules to inside the home cage for quantification of complex operant and feeding behaviours [29], or to connect individual operant test chambers to the home cage by way of a short tunnel [30] to minimise intervention and provide a higher throughput training-testing framework. However, these both have a requirement for animals to remain socially isolated to ensure that the cognitive performance of each individual can be monitored over time. Considering that social isolation itself can induce cognitive deficits [31, 32] and depression-like behaviour [33, 34], this is a huge confound for assessment of cognition in rodent models. In contrast, appropriate social interaction can enhance neuroplasticity [35], emotional and social intelligence [36] and influence performance on complex cognitive tasks [37]. Recently, the capacity to monitor and track rodents in social groups has become achievable with radiofrequency identification (RFID) technology [38, 39] in combination with gating access to test chambers based on a method of automatic animal sorting [28, 40, 41]. The development of a fully automated, experimenter-free method for touchscreen-based cognitive testing in rats has been ongoing since the first prototype was constructed in 2017, allowing the successful adaptation of the trial-unique non-matching to location (TUNL) task in an environment that both eliminates experimenter intervention and allows animals to live in social groups throughout testing [27]. This study demonstrated that the learning rate of self-motivated and undisturbed rats was much faster when experimenter involvement is removed.

The potential to more rapidly test cognition in rodents without experimenter intervention and in social groups opens the door to examine whether cognitive inflexibility predisposes individuals to pathological weight loss in ABA – particularly important because the ABA model develops differently in adult compared to adolescent ages [42]. It also allows us to determine the persistence of inflexibility following weight recovery in ABA rats, in order to use this model to screen novel pharmacotherapeutics for AN. In the present study, we used the automated and experimenter free touchscreen testing system developed from the prototype mentioned above (and now commercially available from PhenoSys, GmbH) to investigate both of these ideas. This automated approach also enables animals to express a more naturalistic behavioural repertoire, a feature ideally suited to comprehensive interrogation with unbiased machine learning approaches to quantify behavioural profiles. Here, we exploited this union with analysis of uninterrupted video recordings of touchscreen sessions using DeepLabCut [43] and B-SOiD [44] to determine the behavioural drivers of cognitive performance. Moreover, the high-throughput pipeline for video analysis from touchscreen sessions that we have established is available openly and may prove useful for future experiments aimed at identifying the behavioural correlates of cognitive performance in rodent models.

## Materials and methods

### Animals and housing

All animals were obtained from the Monash Animal Research Platform, Clayton, VIC, Australia. Initial exploration and optimisation of the novel touchscreen testing system was performed in a cohort of female Sprague-Dawley rats (*n*=20), 6-7 weeks old at the commencement of testing (see **Supplementary Information** for details). To assess both ABA and cognitive behaviour in the same animals, female Sprague-Dawley rats were 5-6 weeks of age upon arrival in the laboratory. Animals were group-housed and acclimated to the 12h light/dark cycle (lights off at 1100h) for 7 days before experiments commenced. To examine whether cognitive flexibility predicted pathological weight loss in ABA, rats (*n*=36) were tested on the pairwise discrimination and reversal learning task and subsequently exposed to the ABA paradigm. To determine whether exposure to ABA altered cognitive performance on the same task, rats (*n*=24) were exposed to the ABA paradigm and allowed to recover to ≥100% body weight prior to pairwise discrimination and reversal learning (see **Fig 1** for timeline of experiments). Rats in each experiment were age-matched for the initiation of ABA exposure, in order to control for age-related changes in vulnerability to weight loss [42].

**Figure 1.**
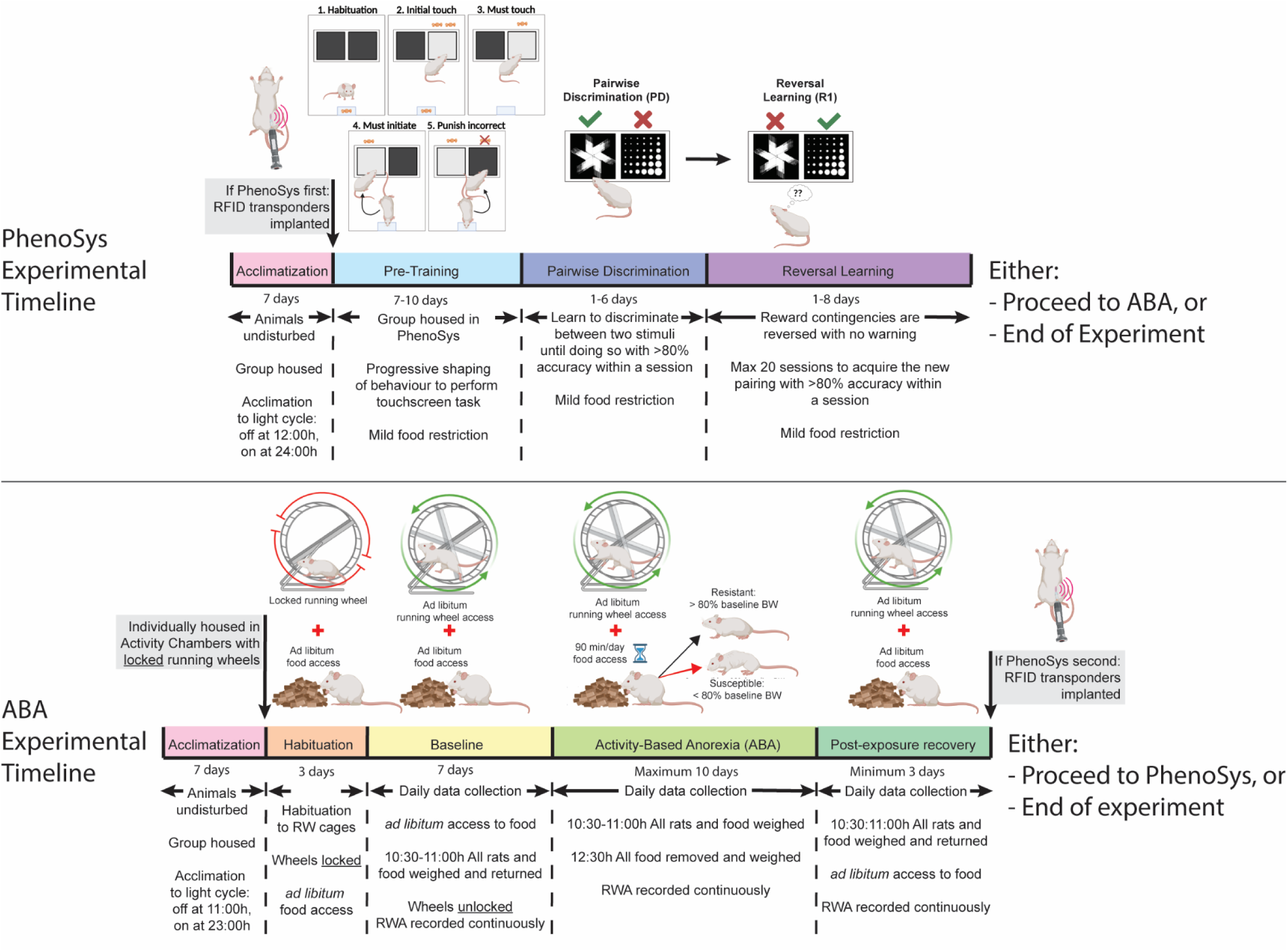
Timeline of experiments. Rats were <u>acclimated</u> to the reversed light cycle for 7 days before each experiment commenced. In experiment 1, rats underwent touchscreen cognitive testing before undergoing the ABA paradigm. In experiment 2, rats were exposed to the ABA paradigm prior to cognitive testing in the PhenoSys apparatus. The PhenoSys cognitive testing paradigm consisted of 7-10 days of <u>pre-training</u>, 1-6 days of <u>pairwise discrimination</u> and 1-8 days of <u>reversal learning</u>. The ABA paradigm consisted of 3 days of <u>habituation</u> with ad libitum food access, 7 days of <u>baseline</u> testing with *ad libitum* access to food and a maximum of 10 days of ABA conditions with time-limited food access (90 min/day) and unrestricted running wheel access, followed by a minimum of 3 days of body weight <u>recovery</u> with reinstatement of *ad libitum* access to food.

A male rat was singly housed in all experimental rooms to synchronise the oestrous cycles of the female rats, known as the Whitten Effect [45]. All experimental procedures were conducted in accordance with the Australian Code for the care and use of animals for scientific purposes and approved by the Monash Animal Resource Platform Ethics Committee (ERM 29143 and 15171).

### Automated sorting and touchscreen testing using PhenoSys

#### Surgical implantation of radio-frequency identification (RFID) transponder

Rats were anaesthetised with isoflurane in oxygen (5% for initiation, 2.5% for maintenance) and subcutaneously implanted with RFID transponders (2.1 x 12mm; PhenoSys, Berlin) into the left flank using a custom designed syringe applicator. The incision site was sealed by tissue adhesive (Vetbond 3M; NSW, Australia).

#### Multimodal apparatus

Following RFID implantation, rats were group housed (*n*=6 per group) in separate home cages of the Phenosys apparatus (**Supp Fig 1**; PhenoSys, Berlin) and allowed to habituate to the home cage with *ad libitum* food access for 1 day prior to behavioural intervention. Food (standard laboratory rodent chow; Barastoc Feeds, AU) was provided daily prior to the dark phase throughout the duration of the experiment to maintain ∼90% of free-feeding body weight. Because of the young age of the animals, this 90% was increased each week by 10% to account for the normal growth curve during development (Charles River Laboratories). The system was housed in a temperature (22-24°C) and humidity (30-50%) controlled room under a reversed 12h light/dark cycle (lights off at 1200h).

This custom designed home cage (26cm x 34cm x 55cm) was placed above an array of twenty RFID readers to track movement of rats. An automated sorter cage connected the home cage to the testing chamber via two plastic tunnels (8.5cm in diameter). The automated sorting system consisted of a sorter cage which was positioned directly above a scale for body weight recording, two RFID readers for animal identification and two software-controlled gates. Selective passage of a single rat from home cage to testing chamber required RFID detection and matching recorded body weight with pre-set weight defined within the PhenoSoft software. The trapezoid testing chamber consisted of two walls, a touchscreen and on the opposing wall a food magazine with an LED light. A test-specific touchscreen mask with windows at the top and bottom that was dependent on the testing/training phase was placed in front of the touchscreen. The touchscreen illuminated with white light through the windows at the top of the mask to act as house light to signal incorrect responses. Sucrose pellets (20mg; Able Scientific, WA, Australia) were used as rewards and delivered from an automated pellet dispenser positioned outside the testing chamber into the food magazine. Touches to the screen and the delivery and collection of food reward were detected by the breakage of infrared (IR) beams. Conditioned reinforcing stimuli consisted of a positive (high) tone and the illumination of LED light within the food magazine. Negative reinforcers involved a negative (low) tone and the house light mentioned above, followed by a “time out” period. Rats were allowed to return to the home cage via the sorter following completion of cognitive tests, which were operated by the PhenoSoft program (PhenoSys, Berlin).

#### Pre-training to shape reward-based behaviours

A series of pre-training stages including Habituation, Initial Touch, Must Touch, Must Initiate and Punish Incorrect were used to shape reward-based behaviours of the rats toward the touchscreen (**Fig 1**). A mask with three side-by-side windows was used in all pre-training stages. Rats were allowed to have multiple sessions of training per day, with a maximum duration of 30 minutes or 30 trials per session and a 1-hour time out period between sessions.

#### Pairwise discrimination and reversal learning

The pairwise discrimination and reversal learning task was used to assess cognitive flexibility in rats. A touchscreen mask with two side-by-side windows was used in the task. Rats were allowed to perform multiple sessions per day, as in pre-training stages, and were first required to discriminate between two stimulus images (**Supp Fig 2E**) and associate touching one of the images with receiving reward. Rats were required to complete 2 sessions (2x30 positive trials) with accuracy >80% within one day to reach progression criterion to reversal learning, in which the stimulus-reward association was reversed. The progression criterion in reversal learning remained the same as in pairwise discrimination. The training and testing protocols were adapted from previous studies [46, 47] with modifications to accommodate to the automated system (**Supp Fig 2; Supp Table 1**). To assess ABA and flexible learning in the same animals, each rat was restricted to a maximum of 20 sessions of reversal learning to prevent touchscreen overtraining. Once rats reached either the progression criterion (i.e. learned the task) or 20 sessions of reversal learning (i.e. did not learn the task), they were either transferred to the running wheel (RW) cages to undergo the ABA paradigm or removed from the experiments and euthanised with 300mg/kg sodium pentobarbitone (Lethabarb; Virbac, Australia) (see **Fig 1**).

### Activity-based anorexia (ABA)

The ABA paradigm used in this experiment consisted of unlimited access to a RW and time-restricted food access. At seven weeks of age, or after reaching the progression criterion of reversal learning, rats were individually housed in transparent living chambers with a removable food basket and a RW (Lafayette Instruments, IN, USA) in a temperature (22-24°C) and humidity (30-50%) controlled room under a reversed 12h light/dark cycle (lights off at 1100h). Rats were allowed to habituate to the living chamber with *ad libitum* food access for 3 days and habituate to the RW for seven days to determine baseline running wheel activity (RWA). RWA was recorded by the Scurry Activity Wheel Software (Lafayette Instruments, IN, USA). During ABA, food access was restricted to 90 minutes per day at the onset of the dark phase (1100-1230h). RWA in the hour before the feeding window (1000-1100h) was considered as food anticipatory activity (FAA). Time-restricted food access persisted for a maximum of 10 days or until rats reached <80% of baseline body weight (ABA criterion). Rats were then allowed to recover to baseline body weight before progression to subsequent cognitive testing or removal from the experiment and euthanised with 300mg/kg sodium pentobarbitone (Lethabarb; Virbac, Australia) (**Fig 1**).

### Machine learning tools for tracking rats

To track the body parts of rats over time, videos in the touchscreen chamber were imported into DeepLabCut (version 2.2.1.1) [43, 48] (https://github.com/DeepLabCut/DeepLabCut). One experimenter labelled 1182 frames from 9 videos with the most variation in camera lighting. We trained a ResNet-50 neural network [49, 50] for 200,000 iterations using a training fraction of 80%. We used 1 shuffle and the errors for test and training were 3.97 pixels and 3.13 pixels respectively. For comparison, the image sizes were 576 by 432 pixels. All default settings were used except a global scale of 1 and *p*-cutoff of 0. See **Supplementary Methods** for details of zone analysis.

To track the behaviours of rats over time, DeepLabCut-tracking data was imported into B-SOiD (version 2.0) [44] (https://github.com/YttriLab/B-SOID). The tracking data for the nose point, left ear, right ear, left hip, right hip and tail base was used to train an unsupervised behavioural segmentation model. The video frame rate was selected as 30 fps. We randomly selected 49% of data and B-SOiD randomly subsampled 12% of that data (input training fraction of 0.12). The minimum time length for clusters to exist within the training data was adjusted to yield 34 clusters (cluster range of 0.17%-2.5%). These 34 clusters were manually grouped into 6 behaviours by interpreting video snippets of behaviours that last >300-ms (see **Supplementary Methods**). These behaviours are grooming, inactive, investigating (nose interacts with either the pellet dispenser or images), locomote (walking forwards), rearing and rotating body. Fleeting behavioural bouts that lasted <300ms were also replaced with the last known behaviour.

The codes used for each of these steps can be found here https://github.com/Foldi-Lab/PhenoSys-data. This includes all the codes and example data needed to reproduce this analysis from the touchscreen chamber videos to the spider plots and time bin heatmaps.

### Exclusions

To assess whether cognitive flexibility predicted susceptibility to weight loss in ABA, rats that failed to reach progression criterion within 20 sessions of reversal learning (First reversal; R1) were excluded from all behavioural and performance data analyses because their levels of flexible learning were unable to be assigned (*n*=3). In addition, three rats demonstrated abnormal weight loss trajectory due to food hoarding in ABA and were therefore excluded from all ABA analyses, considering, as this behaviour confounds the generation of the ABA phenotype. Moreover, one rat failed to recover to >80% baseline body weight during exposure to ABA and one rat failed to learn pre-training to shape reward-based behaviour towards touchscreen after prior exposure to ABA. These two animals were excluded from all data analyses, resulting in a final sample size of *n*=22 in the assessment of effect of prior exposure to ABA on cognitive flexibility. All sessions post-criterion or with technical issues were excluded for performance and behavioural analyses.

### Data processing and statistical analyses

Daily data output files from the touchscreen, sorter and activity monitor were processed using our freely available data analysis pipeline (https://github.com/Foldi-Lab/PhenoSys-codes) to provide detailed information about the performance of each rat during their touchscreen sessions. Statistical analyses were performed using GraphPad Prism 9.1.1 (GraphPad Software, San Diego, CA, USA). Statistical significance was considered as *p*<.05 and analyses including Log-rank (Mantel-Cox) test, two-tailed unpaired t-test, linear regression, correlation, one-way and two-way analysis of variance (ANOVA) with Tukey’s or Bonferroni’s post hoc multiple comparisons were used according to number of groups and type of data. Full details of statistical tests performed in these studies can be found in **Supp Table 2**.

### Data and code availability

The data generated in this paper can be found at https://doi.org/10.6084/m9.figshare.21539685. A data analysis pipeline for providing the key data per session can be found at https://github.com/Foldi-Lab/PhenoSys-codes. The codes used to create the pose estimation and behavioural segmentation analysis and figures can also be found at https://github.com/Foldi-Lab/PhenoSys-data.

## Results

### System validation & optimisation

Prior to experiments involving the ABA model, we first conducted a series of experiments to validate and optimise use of the novel testing system in young female rats. We revealed distinct patterns of behaviour at the reversal of reward contingencies (R1; **Supp Fig 3A-C**), and confirmed that while R1 was more difficult to learn than the initial pairwise discrimination (PD), subsequent reversals were progressively easier to learn (**Supp Fig 4A-C)**. The surprising finding from these initial experiments was that the speed of learning serial reversals was driven largely be reduced omissions at the second and third reversals (R2 and R3; **Supp Fig 4D-E**). One plausible contributor to the high number of omitted trials is the time of day, considering that animals can initiate sessions when they are motivated to perform the task as well as if they are simply exploring the touchscreen chamber. Considering that laboratory rats are well-known to be more active in the dark phase, we compared performance between animals who retained unlimited touchscreen access to those that had access restricted to the dark phase (**Supp Fig 4F-K**). Restricting access to the dark phase increased accuracy in PD (*p=.*0371; **Supp Fig 4G**), which was specific for initial learning, with more substantial between-group differences during the first 100 trials (*p=*.0030; **Supp Fig 4H**). Dark-phase restriction also reduced the number of omitted responses during both PD and R1 (**Supp Fig 4I**), however this was not significantly different overall (**Supp Fig 4J**) but rather restricted to the initial stages of discrimination and reversal learning (PD *p=*.0024; R1 *p=.*0332; **Supp Fig 4K**). The reduced variability in responding within the restricted access group throughout serial reversal learning (see **Supp Fig 4F & I**) is likely to be driven by an increase in motivation that is facilitated by restricted access, and although time of day did not appear to systematically alter performance in animals with unrestricted access (**Supp Fig 5**), we adopted this dark phase restricted approach for subsequent experimental cohorts. Importantly, none of our experimental groups differed in their rate of acquisition of the pretraining stages of the touchscreen task (**Supp Fig 6**), ruling out broad spectrum effects of ABA exposure and susceptibility on visual operant learning.

### Reciprocal interactions between ABA exposure and cognitive flexibility

In order to determine whether individual differences in flexible learning could *predict* susceptibility to pathological weight loss in ABA, we tested animals on the reversal learning task prior to exposure to ABA conditions (**Fig 2A**). Our previous ABA studies demonstrate that rats segregate into susceptible and resistant subpopulations and in the present study, 12/22 (55%) rats exposed to ABA conditions were able to maintain body weight above 80% of baseline throughout the 10 days of ABA, therefore being classified as “resistant”. This allowed us to retrospectively compare reversal learning between groups to assess predisposing factors to pathological weight loss. Susceptible and resistant rats differed on all key ABA parameters (i.e. body weight loss trajectory, food intake, RWA) as we have previously published [51, 52] and resistant rats also spent less time moving than susceptible rats during touchscreen sessions (**see Supp Fig 7**). Rats that went on to be susceptible to ABA were able to learn PD at the same rate as rats that went on to be resistant to ABA, as demonstrated by a similar number of days, sessions and trials to reach performance criterion (**Fig 2B-G**). Interestingly, rats that went on to be *resistant* to weight loss in ABA required significantly more sessions at the first reversal (R1) to reach performance criterion (*p*=.0142; **Fig 2C**), and although this did not translate to a significant increase in overall trials required (**Fig 2D**) it related specifically to an increase in non-correct responses (i.e. incorrect + omitted responses, *p*=.0401; **Fig 2F**) and an increased ratio of non-correct responses (*p*=.0182; **Fig 2G**). However, a large proportion of rats with both ABA outcomes demonstrated a similar learning rate in the early perseverative phase of R1 (first 100 trials), suggesting that there exist overlapping subpopulations of susceptible and resistant animals across a spectrum of cognitive flexibility (**Fig 2H**).

**Figure 2.**
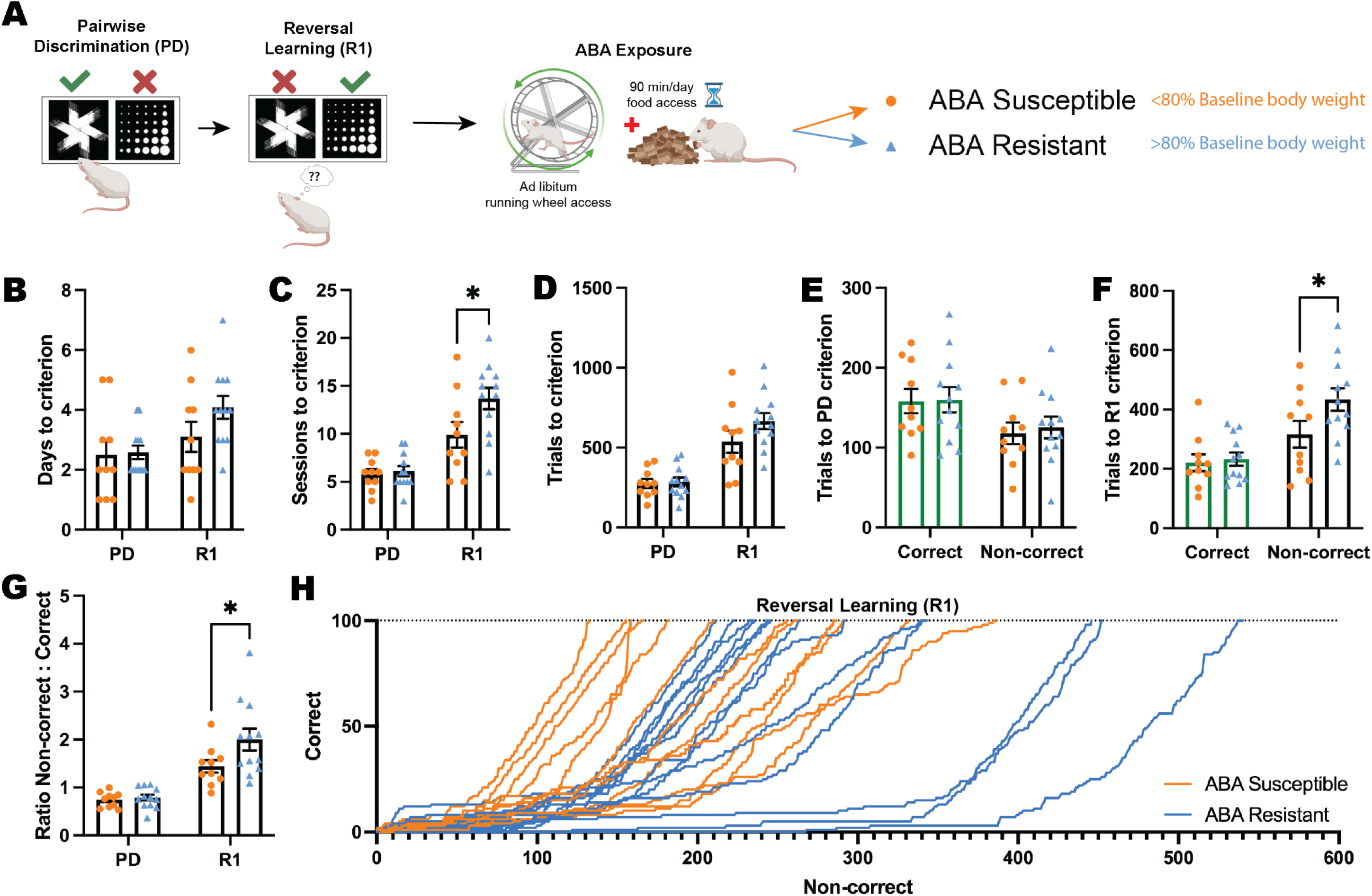
Does cognitive flexibility predict susceptibility to ABA? **(A)** Schematic of pairwise discrimination (PD) and reversal learning (R1) task and subsequent activity-based anorexia paradigm (ABA). Animals split into two experimental groups determined by body weight loss after exposure to ABA: susceptible or resistant to ABA. Bar graphs show group mean ± SEM with individual animals (symbols). **(B)** Number of days to reach criterion. **(C)** Number of sessions to reach criterion (outcome*phase interaction *p*=.0292): R1: ABA resistant > ABA susceptible (*p*=.0142). **(D)** Number of total trials to reach criterion. **(E)** Number of correct or non-correct trials to reach PD criterion. **(F)** Number of correct or non-correct trials to reach R1 criterion (outcome*phase interaction *p*=.0389): Non-correct trials: ABA resistant > ABA susceptible (*p*=.0401). **(G)** Non-correct: correct ratio (outcome *p*=.0399): R1: ABA resistant > ABA susceptible (*p*=.0182). **(H)** Progressive performance across the first correct 100 trials in R1 for individual animals: Non-correct response → X+1; correct response → Y+1. \**p*<.05. For full statistical analysis details and results see **Supplementary Table 2**.

To investigate the behavioural correlates of cognitive task performance that might differentiate rats susceptible versus resistant to weight loss in ABA, we used the DeepLabCut and B-SOiD machine learning tools to annotate videos from touchscreen sessions that were used to train a prediction model, and clustered behaviours based on this model. Analysis of behavioural profiles during touchscreen testing sessions revealed that during initial discrimination learning (PD), rats that went on to be resistant to ABA spent more time engaged in vertical exploration (rearing; *p=*.0336) and locomoting (*p=*.0190) compared to susceptible rats, that also spent significantly more time inactive (*p<*.0001) during touchscreen testing sessions (**Fig 3A**). This differential behavioural profile was similar for reversal learning (R1) sessions, with increased rearing again evident in rats that would go on to be resistant to ABA (*p=*.0384) and increased inactive time for susceptible rats (*p<*.0001), suggesting a consistent exploratory difference between groups even prior to ABA exposure (**Fig 3B**) that may underpin variation in susceptibility to weight loss.

**Figure 3.**
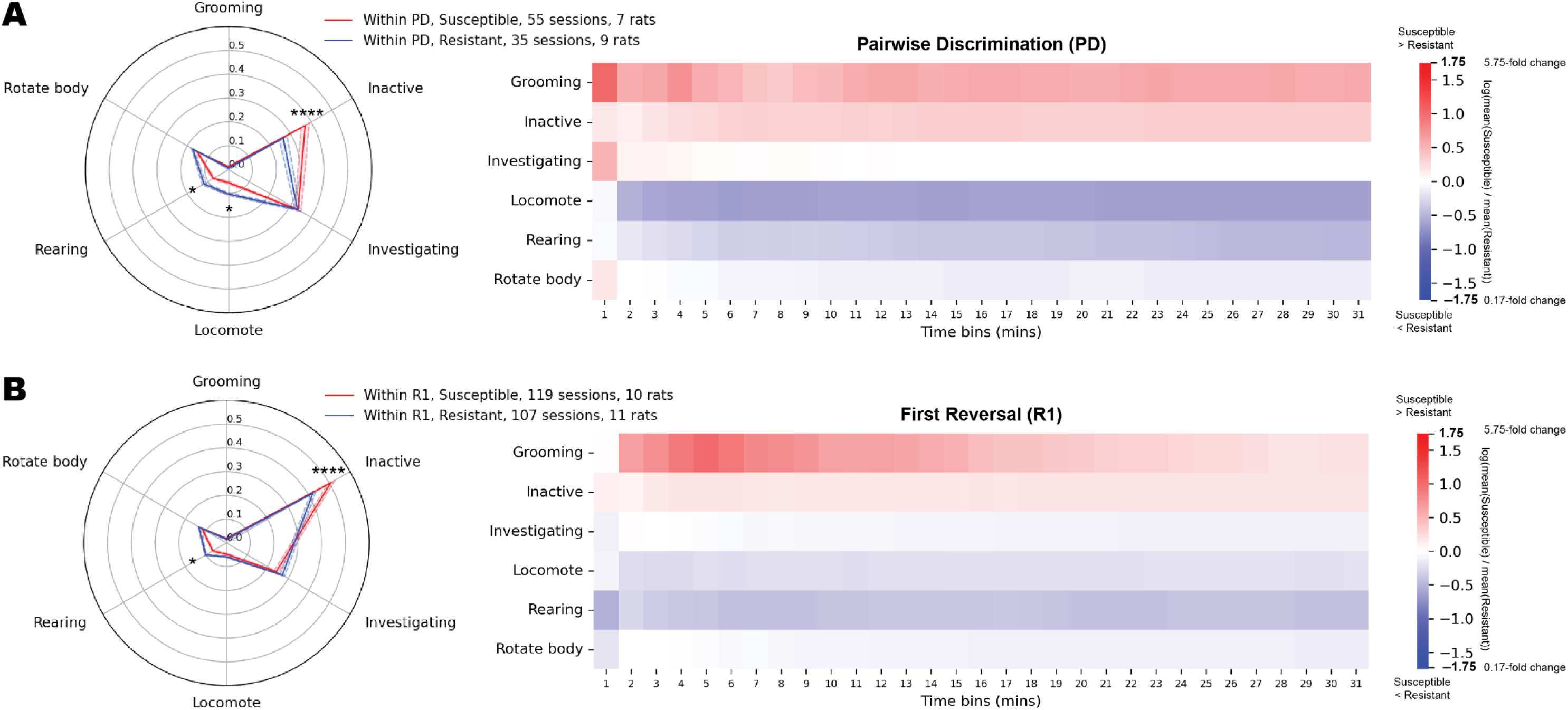
Do behavioural profiles during touchscreen testing sessions predict susceptibility or resistance to ABA? Spider plots and heat maps show the proportion of time spent doing each behaviour within each session video during pairwise discrimination (PD; **A**) or first reversal (R1; **B**). The spider plots show group mean ± SEM (shaded bands). The time bin heat maps show the change in these proportion values between the groups across time. The values are the log(mean(Susceptible)/mean(Resistant)), where log is the natural log. The time bins are cumulative, showing e.g. 0-1 mins, 0-2 mins, etc. **(A)** Within PD (behaviour*outcome interaction *p*<.0001), the susceptible rats spent significantly more time inactive (*p*<.0001) and significantly less time rearing (*p*=.0336) and locomoting (*p*=.0190) than the resistant rats. **(B)** Within R1 (behaviour*outcome interaction *p*<.0001), the susceptible rats spent more time inactive (*p*<.0001) and less time rearing (*p*=.0384) than the resistant rats. \**p*<.05, \*\*\*\**p*<.0001. For full statistical analysis details and results see **Supplementary Table 2**.

To examine whether prior exposure to ABA conditions elicited a persistent change in cognitive flexibility, we allowed animals to recover their body weight to >100% of pre-exposure levels before testing them in the automated touchscreen system (**Fig 4A**). Here, we show that ABA produced a profound impairment in both discrimination and flexible learning, even after weight recovery. Not only were half (50%) of ABA-exposed animals unable to acquire the RL task, compared to 11% of ABA-naïve animals (**Fig 4B**), but those that were able to acquire the task did so at a much slower rate than naïve rats. Exposure to ABA conditions increased the number of sessions required to reach performance criteria (*p=*.0017; **Fig 4C**), however, because the number of sessions animals were allowed each day was capped based on performance, this did not translate to an increased number of days required to reach performance criteria (**Fig 4D**). While the number of trials required to reach PD criteria was not significantly increased for ABA-exposed rats overall (*p=.*0623; **Fig 4E**), the number of correct (*p=*.0231) and omitted (*p=*.0276) trials to acquisition of initial discrimination were higher (**Fig 4F**). In contrast, the number of trials of each type required to learn the reversed contingencies did not differ between ABA exposed and naïve animals that were able to learn the reversal task (**Fig 4G**). Consistent with this, video analysis of touchscreen sessions revealed that during PD, ABA exposed animals spent more time inactive and less time engaged in task-relevant behaviours like rotating, investigating and magazine interactions than did ABA naïve animals, whereas behavioural profiles were more similar between groups for R1 sessions (see **Supp Fig 8**).

**Figure 4.**
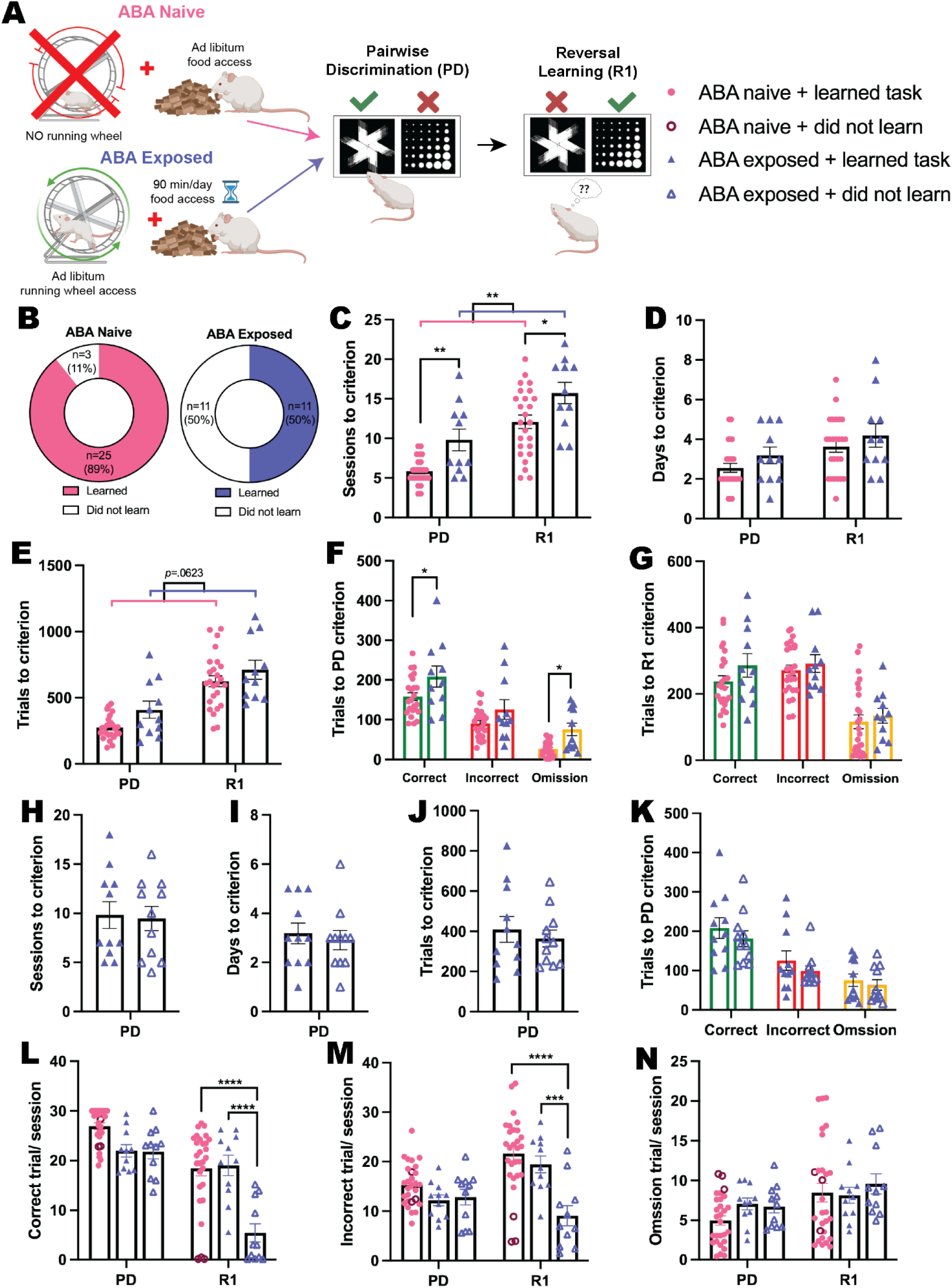
Does exposure to ABA alter cognitive performance? **(A)** Schematic of experimental paradigm showing activity-based anorexia (ABA) Naive or Exposed groups and the subsequent pairwise discrimination (PD) and reversal learning (R1) task. Animals were split into four experimental groups: Naive rats that were not exposed to ABA conditions and learned the reversal learning task (ABA Naive + learned task); ABA Naive but did not learn the task (ABA Naive + did not learn); rats previously exposed to ABA condition that learned the subsequent task (ABA Exposed + learned task); and rats previously exposed to ABA that did not learn the task (ABA Exposed + did not learn). **(B)** Donut plots of experimental groups: 89% (25/28) of the ABA Naive rats learned the reversal learning task compared to only 50% (11/22) of the ABA Exposed rats. **(C)** Number of sessions to reach criterion (exposure *p*=.0017): PD: ABA Exposed + learned task > ABA Naive + learned task (*p*=.0072); R1: ABA Exposed + learned task > ABA Naive + learned task (*p*=.0147). **(D)** Number of days to reach criterion. **(E)** Number of total trials to criterion (outcome *p*=.0623). **(F)** Number of correct, incorrect and omission trials to PD criterion (exposure *p*<.0001): Correct: ABA Exposed + learned task > ABA Naive + learned task (*p*=.0231); Omission: ABA Exposed + learned task > ABA Naive + learned task (*p*=.0276). **(G)** Number of correct, incorrect and omission trials to R1 criterion. Number of **(H)** sessions, **(I)** days, **(J)** total trials and **(K)** correct, incorrect and omission trials to PD criterion. **(L)** Number of correct trials per session (all *ps*<.0003): R1: ABA Exposed + did not learn < ABA Naive and ABA Exposed + learned task (both *ps<*.0001). **(M)** Number of incorrect trials per session (all *ps*<.0030): R1: ABA Exposed + did not learn < ABA Naive (*p<*.0001) and ABA Exposed + learned task (*p=*.0002). **(N)** Number of omission trials per session. Bar graphs show group mean ± SEM with individual animals (symbols). \**p*<.05, \*\**p*<.01, \*\*\**p*<.001, \*\*\*\**p*<.0001. For full statistical analysis details and results see **Supplementary Table 2**.

When considering the response profiles of ABA exposed animals that were unable to learn the reversal task, it was clear that this was not related to impaired performance on aspects of discrimination learning, with similar numbers of sessions (**Fig 4H**), days (**Fig 4I**) and trials (**Fig 4J**) required to reach performance criterion compared to ABA exposed animals that were able to learn. The types of trials required for ABA exposed animals that did and did not learn the reversal task were also unchanged for discrimination learning (**Fig 4K**), however, both the number of correct (*p<* .0001; **Fig 4L**) and incorrect (*p=*.0002; **Fig 4M**) trials per session were substantially reduced for “non-learners” specifically when reward contingencies were reversed (R1). Together with the absence of a significant difference in the number of omitted trials per R1 session (*p>*.9999; **Fig 4N**), this indicates that a lack of reward-based feedback (either positive or negative) impaired the ability of this subgroup of ABA exposed animals to flexibly update responding in the reversal task. The specific impairment in reversal performance was further reflected by more substantial differences between “non-learners” and “learners” in time spent inactive during R1 compared to PD sessions, and by the specific reduction in task-relevant activities including interactions with the reward magazine and touchscreen images in R1 sessions only (see **Supp Fig 9**).

To expore whether cognitive testing changed the development of the ABA phenotype, we compared ABA outcomes in touchscreen-testing naïve animals (Before PhenoSys) to those that occurred following the touchscreen-based reversal learning task (After PhenoSys; **Fig 5A**). Significantly more rats that underwent cognitive testing prior to ABA were able to resist the precipitous weight loss that characterises the model (*p*<.0001; **Fig 5B**) and demonstrated a slow trajectory of body weight loss that plateaued over consecutive days of ABA exposure (**Fig 5C**). When comparing outcomes for both susceptible and resistant animals on key ABA measures, those that had undergone touchscreen testing prior to ABA lost significantly less body weight each day (*p*<.0001; **Fig 5D**), ate more food when food was available (*p=*.0009; **Fig 5E**) and showed a blunted hyperactive phenotype when ABA conditions commenced that was already evident under baseline conditions (*p*<.0001; **Fig 5F**). Although running activity overall was significantly reduced in animals that had previously undergone cognitive testing both at baseline and during ABA (baseline *p=*.0160; ABA *p*<.0001; **Fig 5G**), these rats showed elevated running specifically in the hour preceding food access, known as food anticipatory activity (FAA), which is an adaptive response to scheduled feeding (baseline *p=*.0010; ABA *p*<.0001; **Fig 5H**). While our previous work has shown elevated FAA to be consistently associated with resistance to ABA [17, 51, 52], the increased FAA at baseline for these animals suggests that an anticipatory response was carried over from the scheduled feeding conducted during touchscreen testing. Considering that exposure to cognitive training significantly increased the percentage of rats that were resistant to ABA, it was important to also examine the effects of cognitive training on ABA outcomes in only those rats susceptible to weight loss. The concern was that any differences in ABA outcomes may be driven solely by this subpopulation of resistant animals. Neither mean daily weight loss (**Fig 5I**) nor food intake (**Fig 5J**) were differentially altered by prior cognitive testing in susceptible rats, however, there remained significantly reduced levels of RWA in susceptible rats following discrimination and reversal learning (*p=*.0002; **Fig 5K**). Again, susceptible rats that had previously undergone cognitive training ran less both at baseline (*p=*.0426) and during ABA conditions (*p=*.0165; **Fig 5L**) whereas FAA was specifically increased only during ABA (*p=*.0357; **Fig 5M**). Taken together, these data suggest that cognitive training alters the development of the ABA phenotype specifically through attenuating excessive running activity.

**Figure 5.**
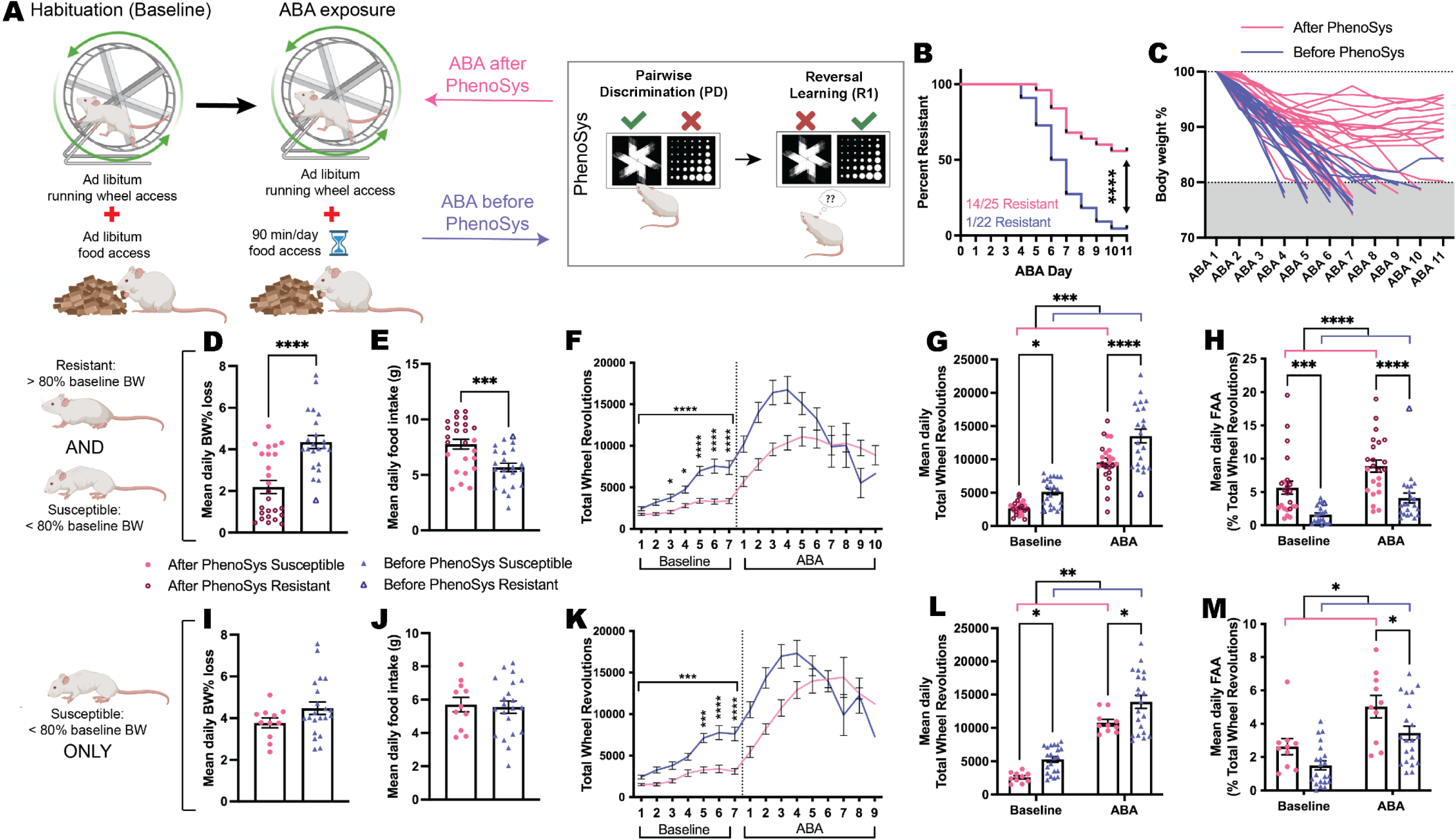
Does cognitive training change the development of the ABA phenotype? **(A)** Schematic of activity-based anorexia (ABA) paradigm and the prior or subsequent pairwise discrimination (PD) and reversal learning (RL) task in PhenoSys. **(B)** Survival plot comparing order effects: ABA resistance was 56% (14/25) for rats that were exposed to ABA <u>after</u> PhenoSys compared to 5% (1/22) for rats that underwent ABA <u>before</u> PhenoSys (*p*<.0001). **(C)** Body weight (% of baseline) trajectories for individual animals across a maximum of 10 days of ABA or until they reached <80%. Data shown are from ALL animals that underwent ABA **(D, E**, **G, H)** or ONLY ABA susceptible animals **(I, J, L, M). (D)** Mean daily ABA body weight (BW) % loss, Before Phenosys > After PhenoSys (*p*<.0001). **(E)** Mean daily ABA food intake, After PhenoSys > Before PhenoSys (*p*=.0009). **(F)** Daily running wheel activity (RWA) across both experimental phases. <u>Baseline</u>, all *p*s<.0001: Before PhenoSys > After PhenoSys (Day 3, *p*=.0440; Day 4, *p*=.0105; Days 5-7, all *ps*<.0001). **(G)** Mean daily RWA (ABA timing *p*=.0002): Before PhenoSys > After PhenoSys during both baseline (*p*=.0160) and ABA (*p*<.0001). **(H)** Mean daily food anticipatory activity (FAA; RWA in the hour before food access; ABA timing *p*<.0001). After PhenoSys > Before PhenoSys during both baseline (*p*=.0010) and ABA (*p*<.0001). Mean daily ABA body weight % loss **(I)** and food intake **(J)**. **(K)** Daily RWA across both experimental phases. Baseline, all *p*s<.0002: Susceptible before PhenoSys > Susceptible after PhenoSys (Day 5: *p*=.0001; Days 6-7: *ps*<.0001). **(L)** Mean daily RWA (ABA timing *p*=.0065): Susceptible before PhenoSys > Susceptible after PhenoSys during both baseline (*p*=.0426) and ABA (*p*=.0165). **(M)** Mean daily FAA (ABA timing *p*=.0157). Susceptible after PhenoSys > Susceptible before PhenoSys during ABA (*p*=.0357). Bar graphs show group mean ± SEM with all individual animals (symbols); line graphs show group mean ± SEM. \**p*<.05, \*\**p*<.01, \*\*\**p*<.001, \*\*\*\**p*<.0001. For full statistical analysis details and results see **Supplementary Table 2**.

## Discussion

Here, we present a validation and optimisation of a novel automated and experimenter-free touchscreen testing platform for rats and demonstrate the application of this system for rapid assessment of cognitive flexibility prior and subsequent to exposure to activity-based anorexia. Critically, the rate of learning in the automated system was shown to be 5 times faster (with approximately 10 times higher throughput) than previously reported with conventional touchscreen testing [17]. While the full spectrum of possibilities arising from the use of the modular PhenoSys touchscreen system are still being realized, the increased throughput, requirement for fewer animals and reduced labour time for experimenters represents a major shift in the way these experiments are conducted and analysed. Our observation that the number of omitted trials is reduced (i.e. engagement is higher) when touchscreen access was limited to the dark phase is consistent with the well-established increase in activity [17] and attention [30] that rats exhibit during the dark period. Moreover, the ability to rapidly test cognitive flexibility with the automated touchscreen system allowed us, for the first time, to examine the cognitive profiles of animals prior to exposure to the ABA paradigm while ensuring that rats remained young adults for ABA exposure. In addition, we were able to conduct this assay in socially appropriate groups and without experimenter intervention, increasing reliability of outcomes and removing potential confounds of handling on subsequent ABA phenotypes [23].

Our previous work revealed that activity within a specific neural circuit (extending from medial prefrontal cortex to nucleus accumbens shell) links pathological weight loss in ABA with cognitive inflexibility on this reversal learning task [17] and suggested that inflexibility might be a biomarker for predicting susceptibility to ABA. The results presented here demonstrate that, contrary to our hypothesis, inflexibility does not predispose animals to the ABA phenotype but instead shows rats that went on to be resistant to ABA were slower to learn the reversal task (i.e. were more inflexible) than ABA susceptible rats. This raises the intriguing possibilities that either inflexibility develops coincident with pathological weight loss in the ABA model or that inflexibility is somehow protective against ABA-induced weight loss. One finding supporting the latter is that rats that went onto be resistant to ABA were hyper-exploratory in touchscreen testing sessions, evidenced by increased rearing behaviours and decreased time spent inactive during the task. Regarding the former, while we did not examine flexible learning *during* exposure to ABA conditions, the idea that inflexibility and ABA develop in concert fits with the timing of neural circuit manipulation used in our previous work [17]. That is, both pathological weight loss and inflexibility were prevented by suppressing the same “cognitive control” neural circuit, but suppression occurred *during* the development of ABA, not prior to ABA exposure. Future studies should delineate the precise stage during the development of the ABA phenotype that inflexibility becomes apparent, thereby defining a “therapeutic window” in which novel pharmacological treatments could be tested with greater translational relevance.

Considering a major hallmark of the ABA phenotype is the development of paradoxical hyperactivity when restricted food access is imposed, it has been suggested that animals susceptible to weight loss in ABA are unable to effectively adapt exercise behaviour to the change in food availability. And yet reversal learning, arguably a “cleaner” test of an ability to effectively adapt behaviour to environmentally imposed change, was *improved* in rats that went on to be susceptible to weight loss in ABA in the present study. This challenges our conceptualisation of so-called “compulsive” [53] wheel running that occurs during ABA and precipitates the rapid weight loss characteristic of the model. Even after decades of experimental use of the ABA model, the causes for this paradoxical hyperactivity remain elusive. A recent study in ABA mice demonstrated that a loss of behavioural flexibility following disrupted cholinergic activity in the dorsal striatum was associated with both facilitated habit formation and increased vulnerability to maladaptive eating [54] but neither the accelerated formation of habits or inflexible behaviours were associated with changes in hyperactivity. Similarly, although compulsive behaviour in individuals with AN as been described to develop under more habitual than goal-directed control [55] these associations have been restricted to eating behaviour rather than exercise. Could it be that excessive exercise in ABA rats (and possibly individuals with AN) represents not a compulsion or habitual behaviour but rather a form of extreme goal-directed control? Understanding how wheel running in ABA might be differentiated from habitual or compulsive behaviour would allow us to better probe the neural circuits underlying the development of ABA. The corollary of this would be the potential to improve cognitive-behavioural therapy for patients with AN based on a perspective of modifying eating disorder-relevant goals, particularly in those ∼80% of individuals with AN that engage in excessive exercise [56]. Combining the ABA model with cognitive behavioural assays that contrast habitual with goal directed behaviour, including outcome devaluation tasks [57, 58], could aid substantially in this understanding.

Our data also suggests that operant training prior to exposure to ABA also alters the subsequent development of anorectic phenotypes, particularly by reducing the maladaptive wheel running that typifies the ABA model. While the independent effects of sucrose consumption and scheduled feeding, both procedural aspects required for touchscreen testing, on subsequent weight loss in ABA are yet to be determined, if ABA indeed develops through a failure to effectively adapt to the change in food availability, then our results support the idea that experience with flexible learning tasks improves this adaptive capacity. Interestingly, this aligns with recent evidence that increased cognitive flexibility mediates improvements in eating disorder symptoms in patients with AN [14].

While the identification of a behavioural predictor (or biomarker) for pathological weight loss in ABA remains a challenge, the finding that rats exposed to ABA subsequently showed marked impairments in both discrimination and reversal learning, even after body weight recovery, is entirely in line with the clinical presentation of inflexibility in patients with AN long after body weight recovery [7, 59, 60]. Intriguingly, this learning impairment induced by ABA exposure was evident from the first session of each phase of training and even within the first 10 minutes of initial PD performance (see **Supp Fig 10**). This lends weight to the use of the ABA model as an effective tool with which to probe the biological mechanisms underlying cognitive deficits in AN. Our finding is in contrast to the only other published report of flexible learning after exposure to ABA [61], in which reversal learning was impaired at low body weight in ABA rats but ameliorated with weight recovery. Although the reasons for this discrepancy remain unclear, the touchscreen testing system used in the present study differs on multiple procedural levels from the attentional set-shifting task previously used to examine flexible learning, and our results suggest that the visual reversal learning task may be preferable for delineating the lasting effects of ABA exposure on cognitive function. That we observed impairments following ABA not only on flexible updating of operant responses but also initial discrimination learning points to a potential motivational deficit induced by ABA, in line with our previous work demonstrating a role for the mesolimbic dopamine circuitry in the development and maintenance of the ABA phenotype [62]. Considering that exercise behaviour in AN is also linked with dopaminergic activity [63], future studies should define the time course over which motivation or reward-based deficits arise during ABA and the specific influence of ABA on dopamine signalling in response to reward anticipation and receipt using in vivo fiber photometric recordings paired with detection of dopamine release (using the GRAB_DA sensor; [64]) or dopamine binding (using the dLight sensor [65]).

Not only does the automated touchscreen testing system described here allow us to identify cognitive profiles that more accurately reflect the naturalistic behaviour of animals, but the incorporation of behavioural segmentation using machine learning also assisted with reducing experimenter biases that are commonly found with manual behavioural scoring. The application of DeepLabCut and B-SOiD to the prediction of behaviours in the present study has allowed the manner in which rats complete touchscreen tasks to be determined and revealed a differential behavioural profile during testing for rats that were subsequently susceptible or resistant to weight loss in ABA. Using these tools also enabled the scoring of very large datasets, such as the 185 hours of footage analysed here. Incorporating this type of analysis with animal models that mimic specific aspects of human pathologies will take us closer than ever before to the identification of biological predictors of pathological weight loss in activity-based anorexia that could be used in the early detection of anorexia nervosa in at risk individuals.

## Acknowledgements

The authors acknowledge the incredible technical support from Dr Karsten Krepinsky (PhenoSys, GmbH; Berlin) and the use of Biorender.com in the generation of some figures. We are grateful for financial support for these studies from the Rebecca L. Cooper Medical Research Foundation (Project Grant PG2019373-Foldi) and the National Health and Medical Research Council of Australia (Ideas Grant GNT2001722-Foldi).

## Supplementary Methods

**(A-D):** Pre-processing, pose estimation, zone analysis and behavioural clustering of rats in the touchscreen chamber. (A) FFMPEG was used to pre-process the videos before analysis (https://github.com/FFmpeg/FFmpeg). (B) DeepLabCut was used to predict the locations of the rat body parts (https://github.com/DeepLabCut/DeepLabCut). (C) DLCAnalyzer was to find the time spent in zones (https://github.com/ETHZ-INS/DLCAnalyzer). (D) B-SOiD was used to find the time spent doing different behaviours (https://github.com/YttriLab/B-SOID).

### A Processing videos before they were analysed

1. Videos of the PhenoSys touchscreen chamber were recorded for each experiment (which lasted ∼24 hours). The cameras were infrared, the angle was bird’s eye view, the resolution was 960x720, .mov was the file format and the Multicam software was used to record videos.
2. These ∼24 hour videos were snipped into videos of individual sessions (up to 31 mins). The session start and end times were found from the raw data file generated by the PhenoSys touchscreen system. The start of the video was calibrated to the “start” experiment event in the raw data file. 30 secs before the session start time and 30 secs after the end time were also included in the snipped videos.
3. A few videos that have 1280x720 dimensions were cropped to 960x720, blurry videos were sharpened, all videos were downscaled to 576x432 and converted to the .mp4 file format. Videos that had black frames, frozen frames, where the mouse was completely absent, the pellet magazine was covered, there was high glare or the video was corrupted were excluded.

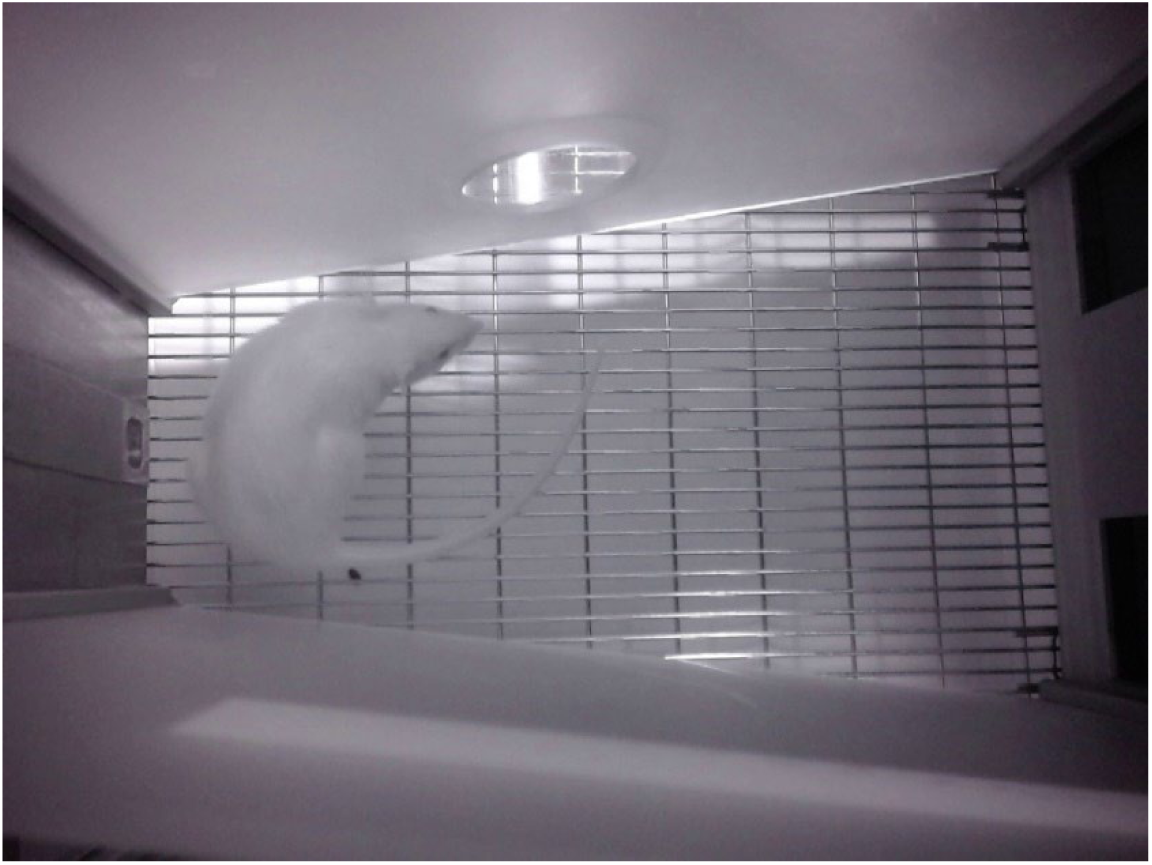

### B Predicting body part locations over time

1. The videos of rats in the touchscreen chamber of the PhenoSys system were imported into DeepLabCut (version 2.2.1.1).
2. One individual labelled 1182 frames from 9 videos with the most variation in camera lighting. The following body parts were labelled:

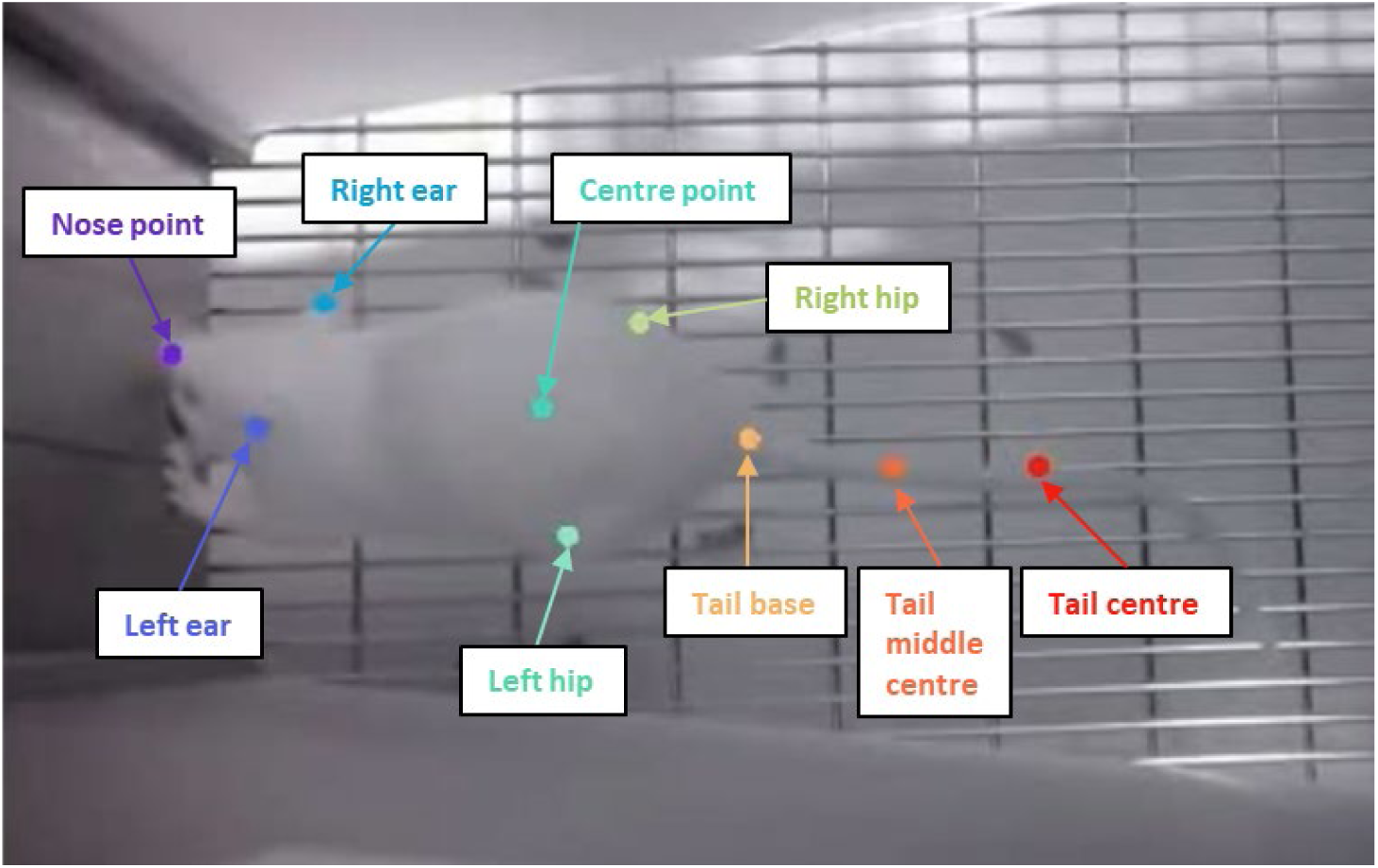
3. We trained a ResNet-50 neural network for 200,000 iterations using a training fraction of 80% and a shuffle of 1.

- The errors for test and training were 3.97 pixels and 3.13 pixels respectively (where the image sizes were 576x432 pixels).
- All default settings were used except a global scale of 1 and a *p*-cutoff of 0.

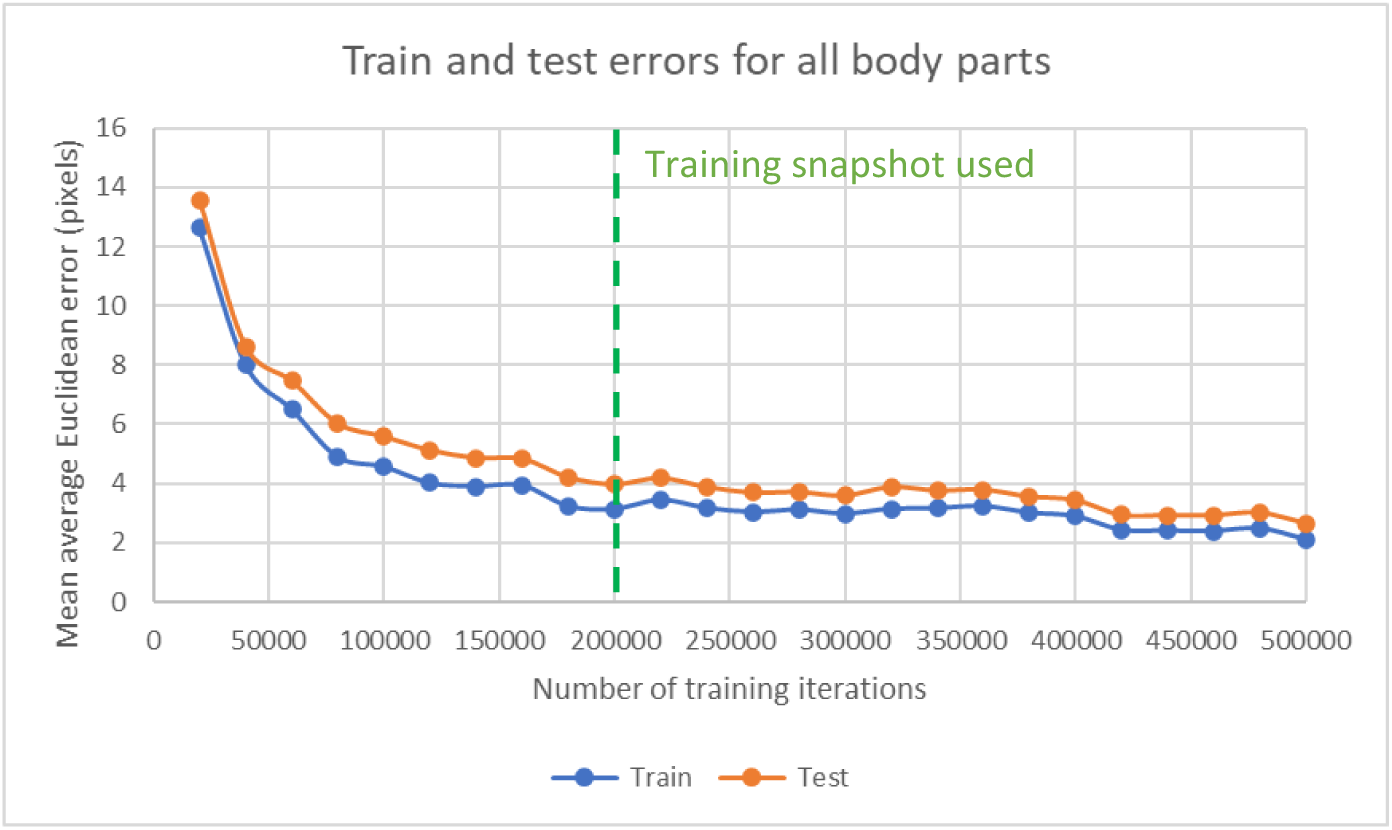

### C Finding time spent in zones

1. All DeepLabCut predictions for the nose point were smoothed using a median filter with a window duration of 0.17 secs (5 frames).
2. The following zones were manually drawn. Distances were calibrated from pixels to cm using the width of the touchscreen wall at 22 cm.

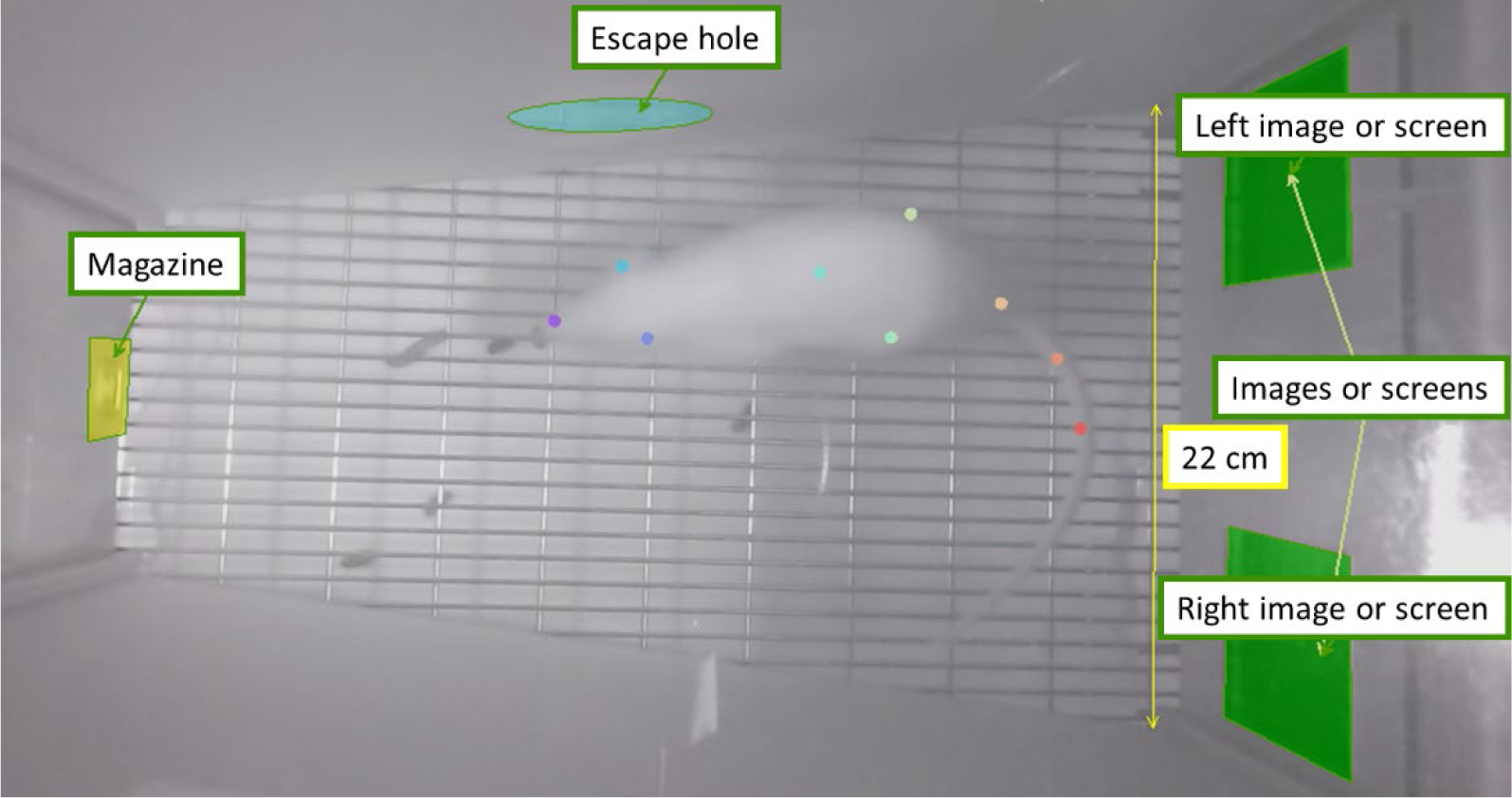
3. Exits out of the arena were defined as the time points when the centre-point prediction *p*-values < 0.05. Here is a characteristic video with the time points highlighted when the rat leaves the touchscreen chamber.
4. This data was imported into DLCAnalyzer. The speed and acceleration were calculated by integrating the nose position over time. A movement cut-off of 5 cm/s was used as the minimum speed to be considered moving. The time spent in each zone was calculated using an integration period of 0.17 secs (5 frames). This defines the minimum time period for a zone transition to occur.

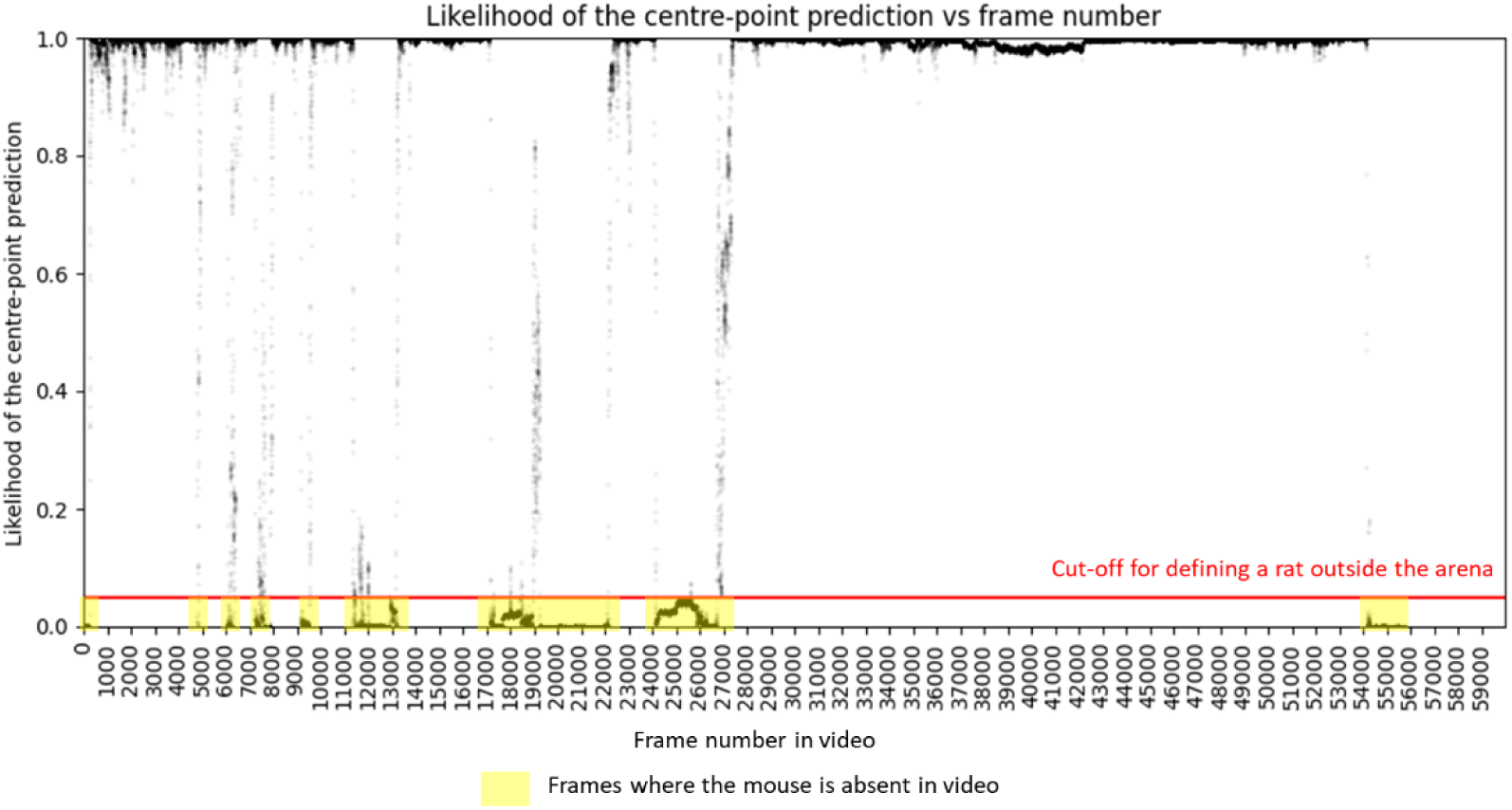

### D Finding time spent doing behaviours

1. The unfiltered DeepLabCut predictions for nose point, left ear, right ear, let hip, right hip and tail base were imported into B-SOiD (Version 2.0). The video frame rate was selected as 30 fps. We randomly selected 49% of all data and B-SOiD randomly subsampled 12% of that data (input training fraction of 0.12).
2. B-SOiD uses UMAP to transform the higher-dimensional pose data into a lower-dimensional space and finds clusters using HDBSCAN. The minimum time length for clusters to exist was adjusted to yield 34 clusters (cluster range of 0.17%-2.5%). These clustered features are then used to train a random forests (RF) classifier.
3. We evaluated our model by examining the UMAP plot, using a box-plot and normalised confusion matrix.

- The UMAP plot shows the prediction of clusters is not significantly dominated by any one given cluster.
- The boxplot below shows the high performance (mean of 0.96) of the random forests classifier on 20% of the training data.
- The normalised confusion matrix shows the high number of true positives (close to 1) and low number of false positives (close to 0) for each behaviour from 0-33 in the training data.

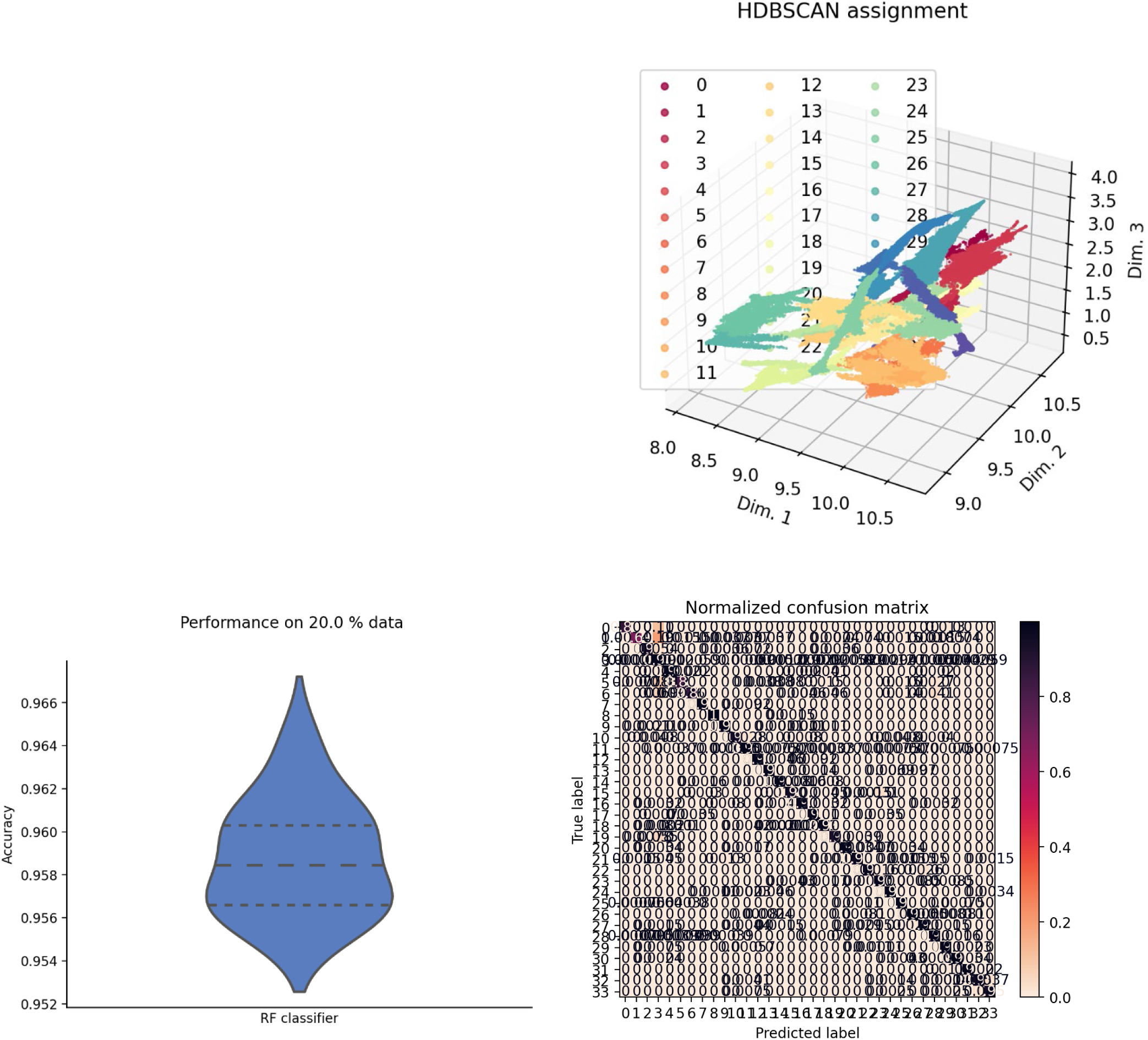
4. These 34 clusters were manually grouped into 6 behaviours by interpreting video snippets of behaviours that last > 300 ms (see supplementary video 1 at https://doi.org/10.6084/m9.figshare.21556677.v1). These behaviours are grooming, inactive, investigating, locomote, rearing and rotate body.

5. Behaviours that lasted < 300 ms were not accurate to the behaviour type. Thus, fleeting bouts that lasted < 300 ms were replaced with the last known behaviour (see supplementary video 2 at https://doi.org/10.6084/m9.figshare.21556677.v1).

| Behaviour | Cluster number | Description of behaviour |
| --- | --- | --- |
| Grooming | 1, 2, 4, 5, 19 | Nose rubbing left/right side of body, left/right paw scratching the face |
| Inactive | 0, 3, 6, 7, 8, 9, 10, 12, 13, 14, 15, 16, 17, 18, 20, 21, 23, 24 | Sitting completely still or moving slightly |
| Investigating | 11, 22, 26, 27, 31, 32, 33 | Nose interacts with images or pellet dispenser |
| Locomote | 30 | Walking forwards |
| Rearing | 29 | Front paws/head touching the walls or unsupported rearing |
| Rotate body | 25, 28 | Rotate body left/right |

---

**Supplementary Figure 1.**
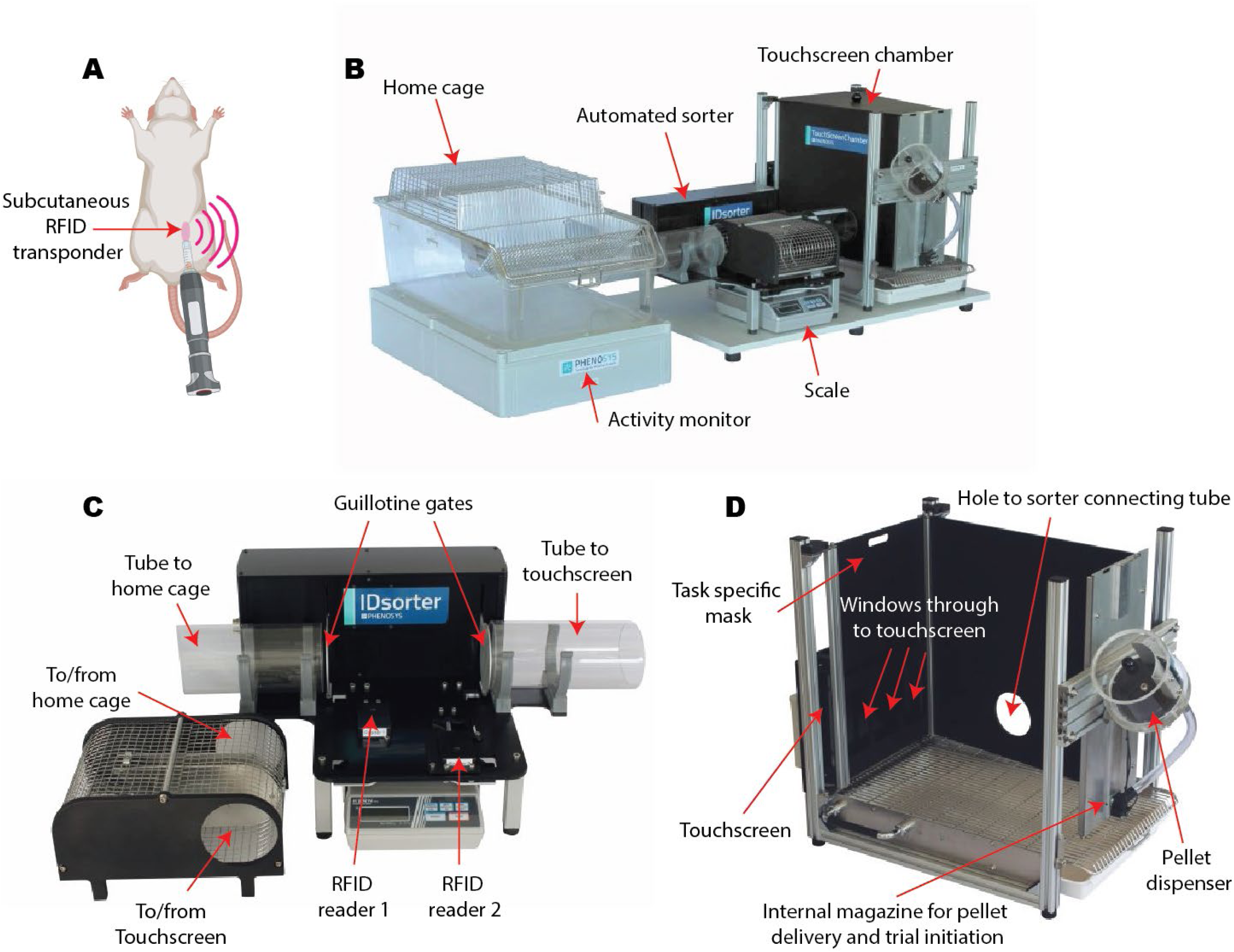
The PhenoSys is an automated home cage and touchscreen testing system. **A)** Animals are implanted with unique radiofrequency identification (RFID) transponders**. B)** Overview of the entire PhenoSys system, in which a home cage is connected to a touchscreen testing chamber via a series of tunnels and a sorting device positioned over a weight scale. **C)** The sorter cage has two RFID readers positioned underneath, and metal guillotine gates control the passage of animals between the home-cage and touchscreen **D)** The touchscreen chamber has an externally mounted pellet dispenser, from which rewards are delivered into a magazine on the opposite side of the chamber from the touchscreen. Images adapted from https://www.phenosys.com/products/touchscreen-chamber/.

**Supplementary Figure 2.**
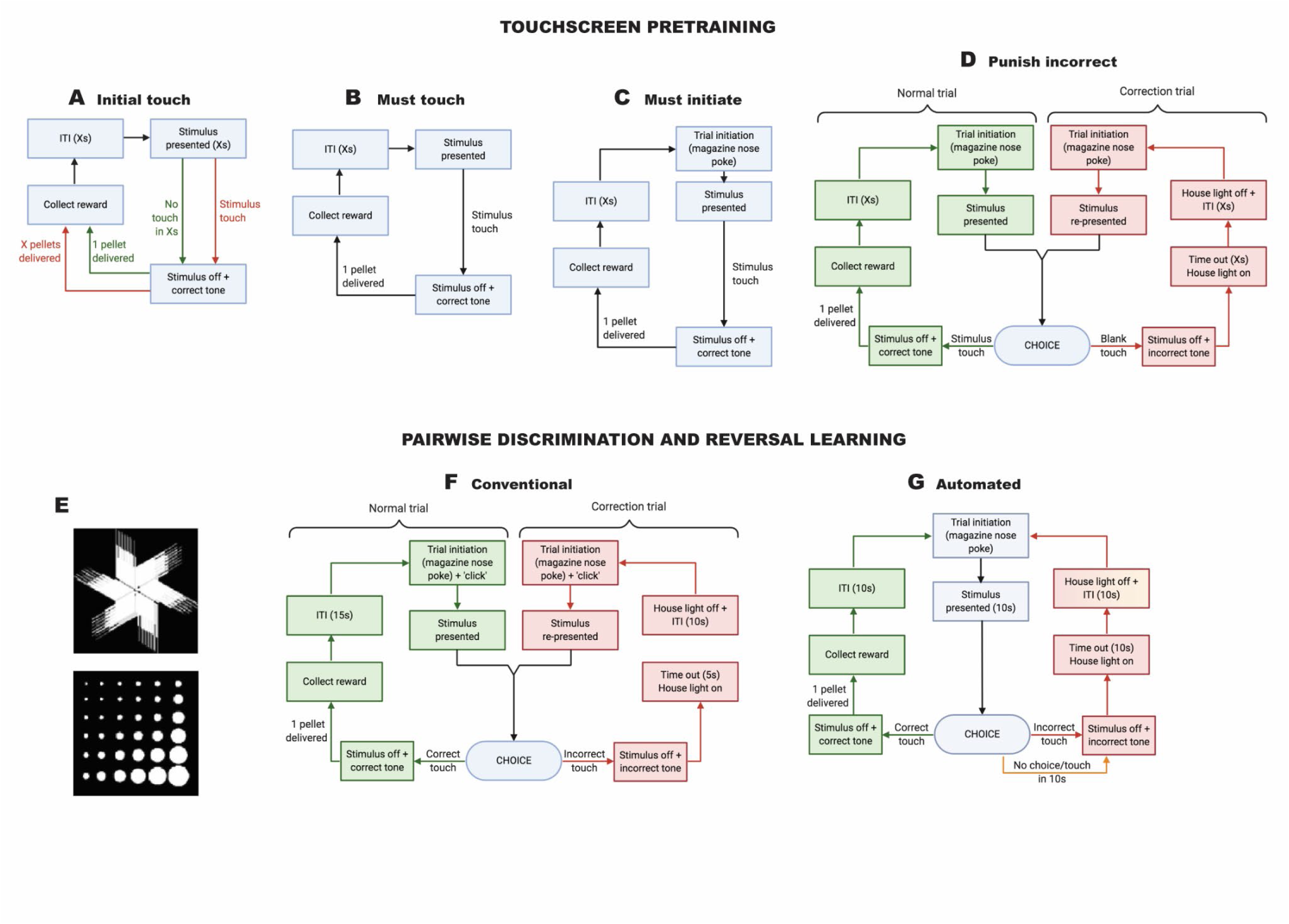
Schematic overview of touchscreen pre-training and serial reversal learning protocol. Image based on [46, 47] and adapted for our protocol. See **Supplementary Table S1** for specific parameters in each stage for PhenoSys touschreen protocol. **E)** The fan/pinwheel (top) and marble array (bottom) images used for pairwise discriminations and all stages of reversal learning in all experiments.

**Supplementary Figure 3.**
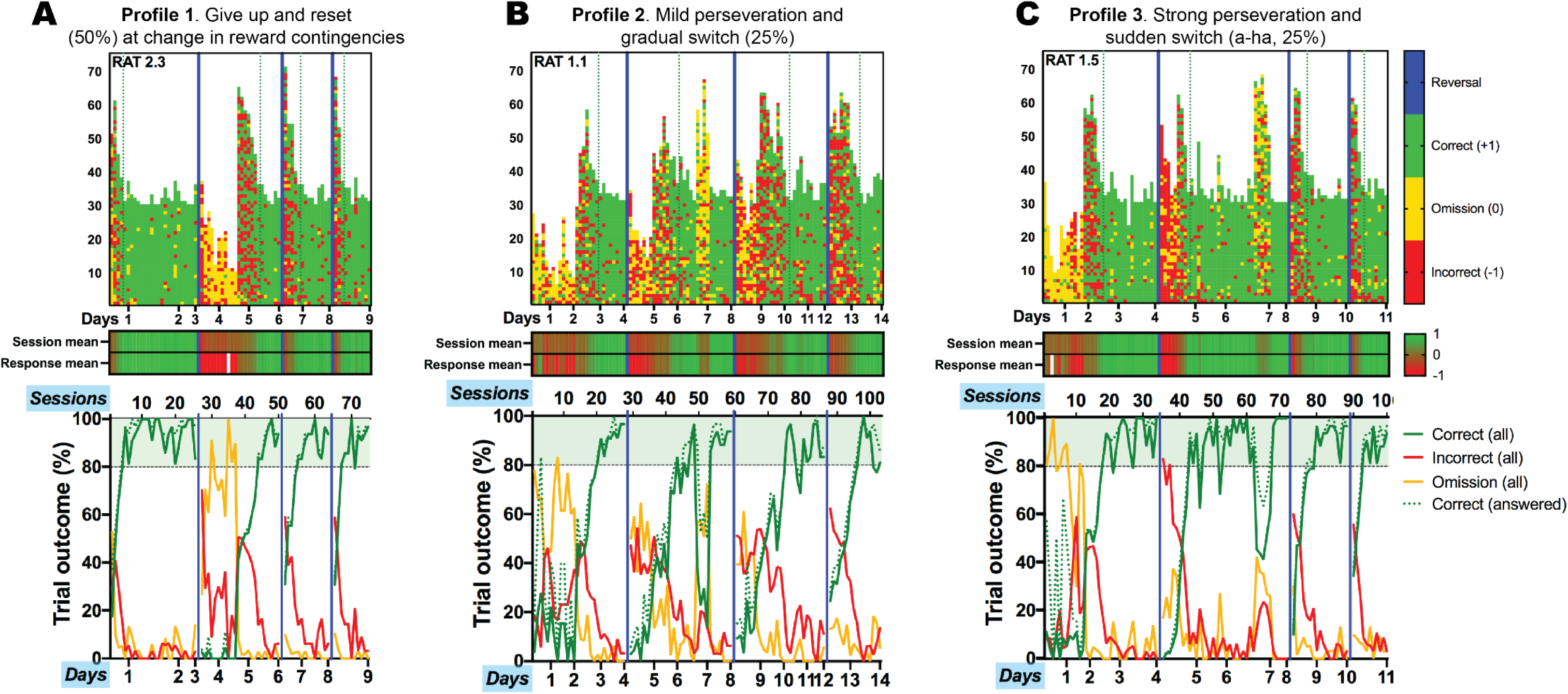
Types of response profiles in female rats using the PhenoSys automated touchscreen system. For initial validation and optimisation of the PhenoSys testing system, female Sprague-Dawley rats (n=20; 6-7 weeks old) were obtained from the Monash Animal Research Platform (Clayton, VIC, Australia) and habituated to the laboratory environment (20-23 °C; 35-65% humidity) under a reverse 12 h light/dark cycle for 7 days prior to testing. Half of these animals were allowed continuous and unlimited access to the touchscreen testing chamber, whereas the other half were allowed access only during the dark phase of the light cycle. Three distinct profiles emerged that were characterised by **(A)** lack of engagement followed by a “reset”, **(B)** mild perseveration and gradual switch, or **(C)** strong perseveration and a sudden switch. **Upper)** Trial-by-trial data with every pixel representing the outcome of a trial and every column representing a single session in chronological order. **Middle)** Average outcome of all trials initiated (session mean) and all trials responded to (response mean) in each session. A summary of performance in each session (each column) was determined by calculating the mean outcome of all trials in a session (correct = +1, omission = 0, incorrect = -1) and the mean outcome of trial responses in a session (i.e. percentage correct; correct = +1, incorrect = -1). **Lower)** Percentage of trial outcomes in all trials within the session (solid lines) and percentage of correct trials for only trials responded to (dotted green line). Blue: reversal of reward contingencies; green: correct trial; yellow: omission trial; red: incorrect trial.

**Supplementary Figure 4.**
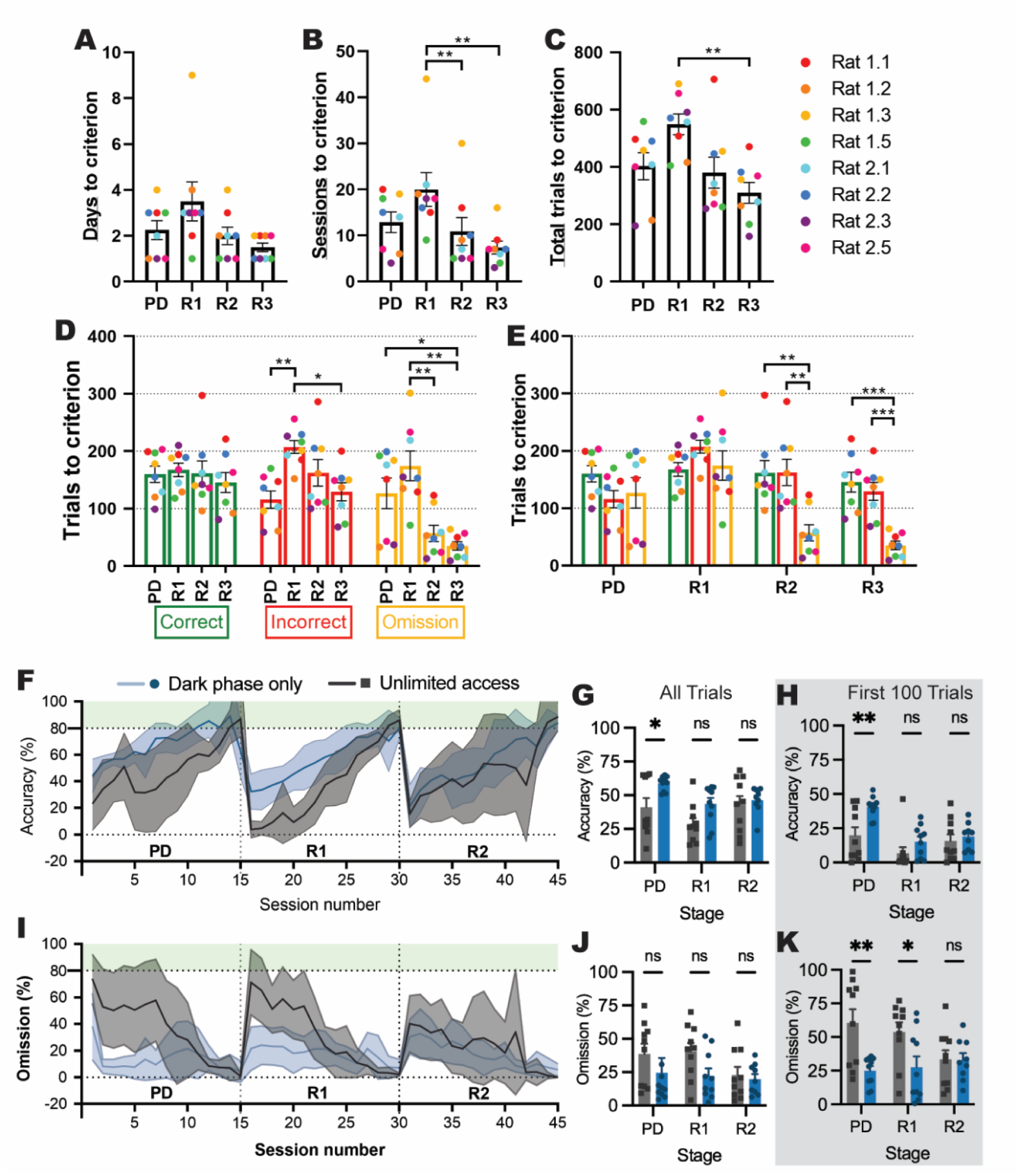
Learning rate over serial reversals and effects of unlimited versus dark-phase only access on cognitive performance in the PhenoSys. **(A-E)** Response types over pairwise discrimination (PD), first reversal (R1), second reversal (R2) and third reversal (R3) with unlimited touchscreen access. **(A)** Number of days to criterion (*p*=.0484). **(B)** Number of sessions to criterion (*p*=.0092): R1>R2 (*p*=.0099), R1>R3 (*p*=.0070). **(C)** Number of total trials to criterion (*p*=.0034): R1>R3 (*p*=.0035). Outcome of trials to criterion grouped by outcome **(D;** *p*=.0032) and by phase **(E;** *p*<.0001). **(D)** Incorrect: R1>PD (*p*=.0014), R1>R3 (*p*=.0309); Omission: PD>R3 (*p*=.0484), R1>R2 (*p*=.0092), R1>R3 (*p*=.0018). **(E)** R2: correct > omission (*p*=.0045), incorrect > omission (*p*=.0059); R3: correct > omission (*p*=.0005), incorrect > omission (*p*=.0008). **(F-K)** Effects of unlimited versus dark-phase only access on cognitive performance. **(F)** Percentage of correct trials across 15 sessions of each phase of the experiment. **(G)** Percentage of correct trials (access *p*=.0039): PD: Dark phase only > unlimited access (*p*=.0371). **(H)** Percentage of correct trials in the first 100 trials (access *p*=.0030): PD: Dark phase only > unlimited access (*p*=.0030). **(I)** Percentage of omission trials across 15 sessions of each phase of the experiment. **(J)** Percentage of omission trials (access *p*=.0737). **(K)** Percentage of omission trials in the first 100 trials (access *p*=.0008): PD: Dark phase only > unlimited access (*p*=.0024); R1: Dark phase only > unlimited access (*p*=.0332). Bar graphs show group mean ± SEM with individual animals (symbols). Line graphs show group mean ± SEM (shaded bands). \**p*<.05, \*\**p*<.01, \*\*\**p*<.001. For full statistical analysis details and results see **Supplementary Table 2**.

**Supplementary Figure 5:**
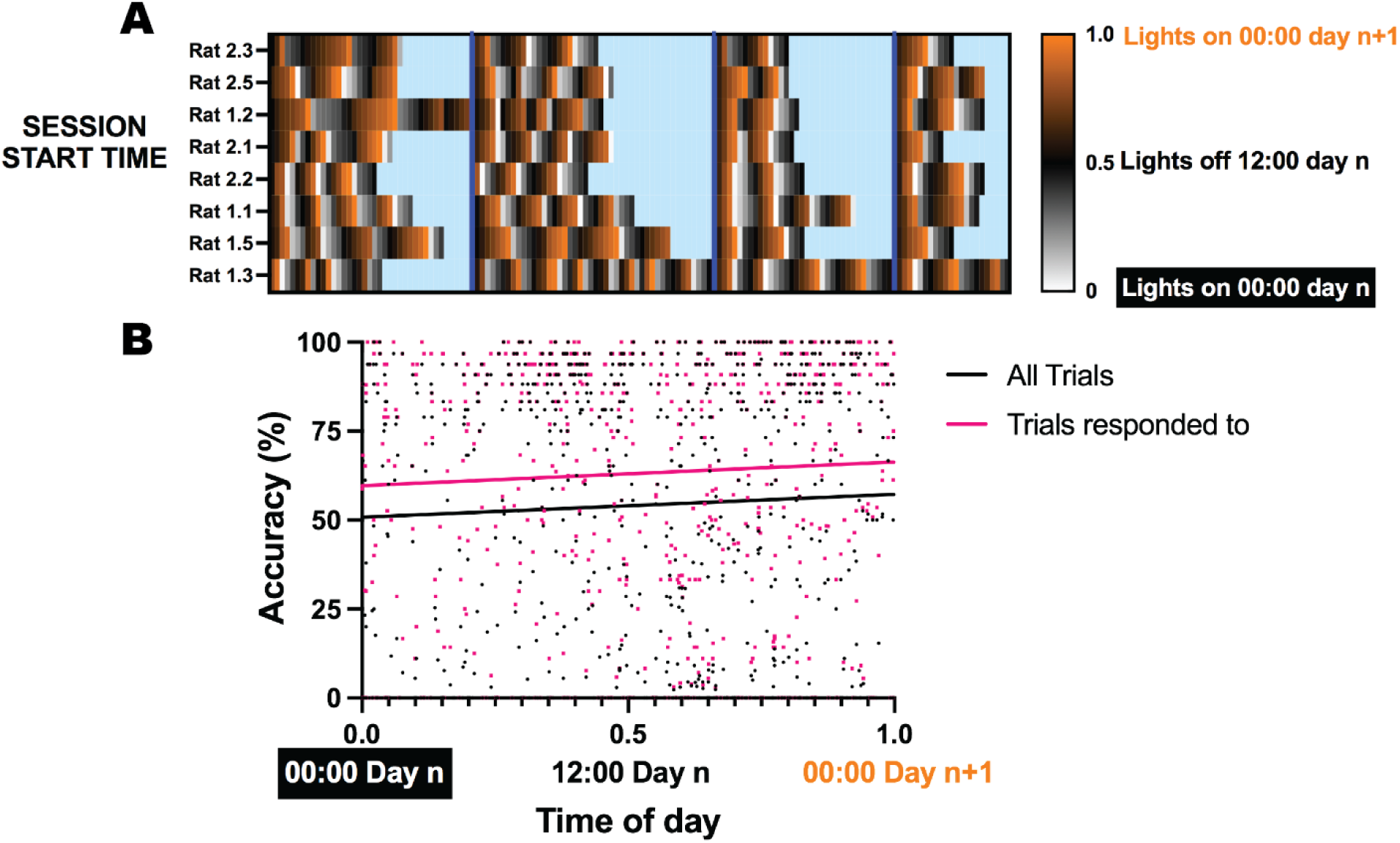
Time of day does not influence PhenoSys touchscreen performance. **A)** Start time of each session. **B)** Correlation between performance and time of day for all trials and only those that elicited a response (**ALL** *r*=.0465, *R*^2^=.0022, *p*=.2088; **RESPONDED** *r*=.0516, *R*^2^=.0027, *p*=.1766).

**Supplementary Figure 6:**
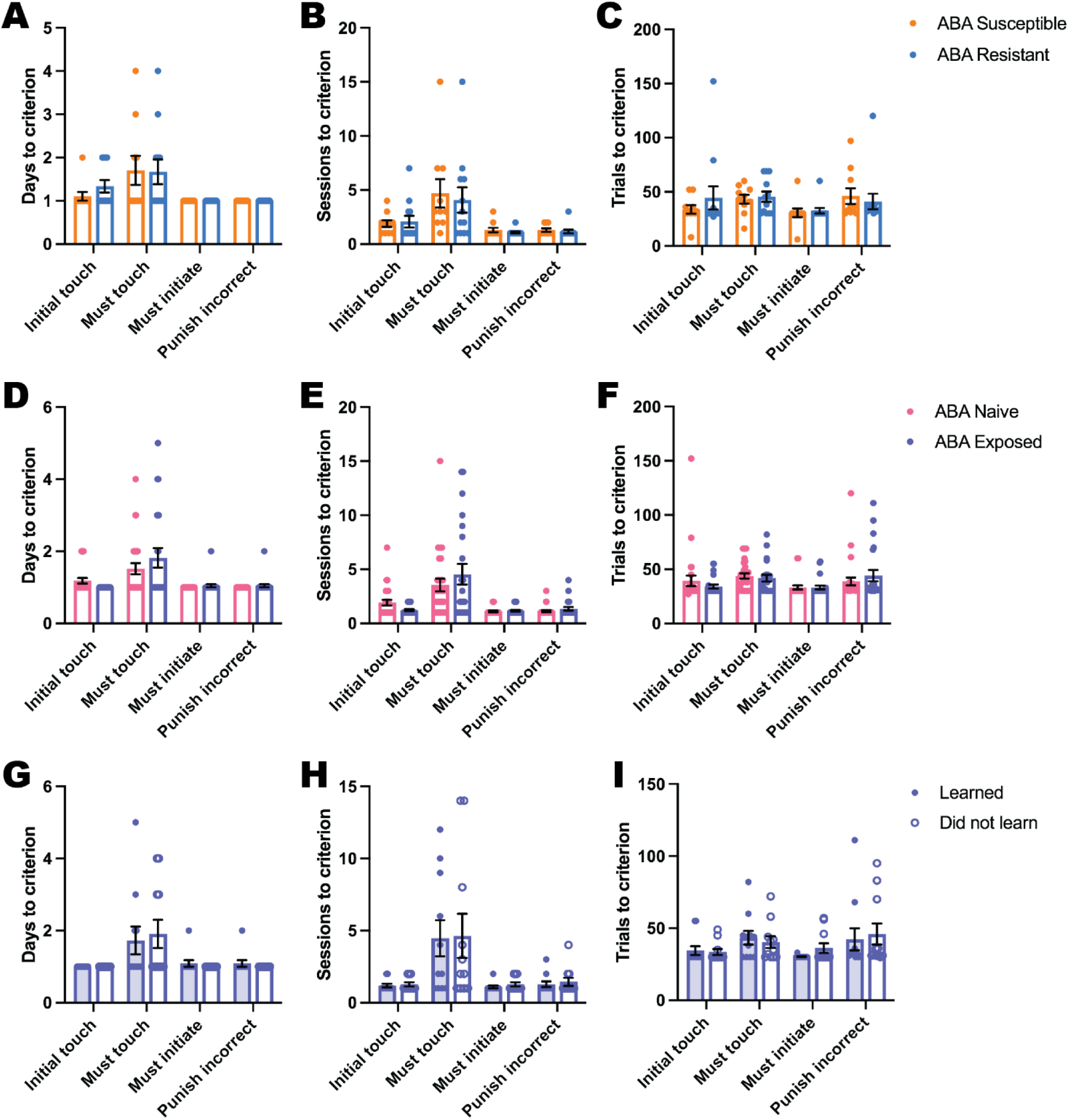
Touchscreen pre-training performance measures. Touchscreen pretraining performance measured by days **(A, D, G)**, sessions **(B, E, H)** and total trials **(C, F, I)** to criterion did not systematically differ between any groups. Bar graphs show group mean ± SEM with individual animals (symbols). For full statistical analysis details and results see **Supplementary Table 2**.

**Supplementary Figure 7:**
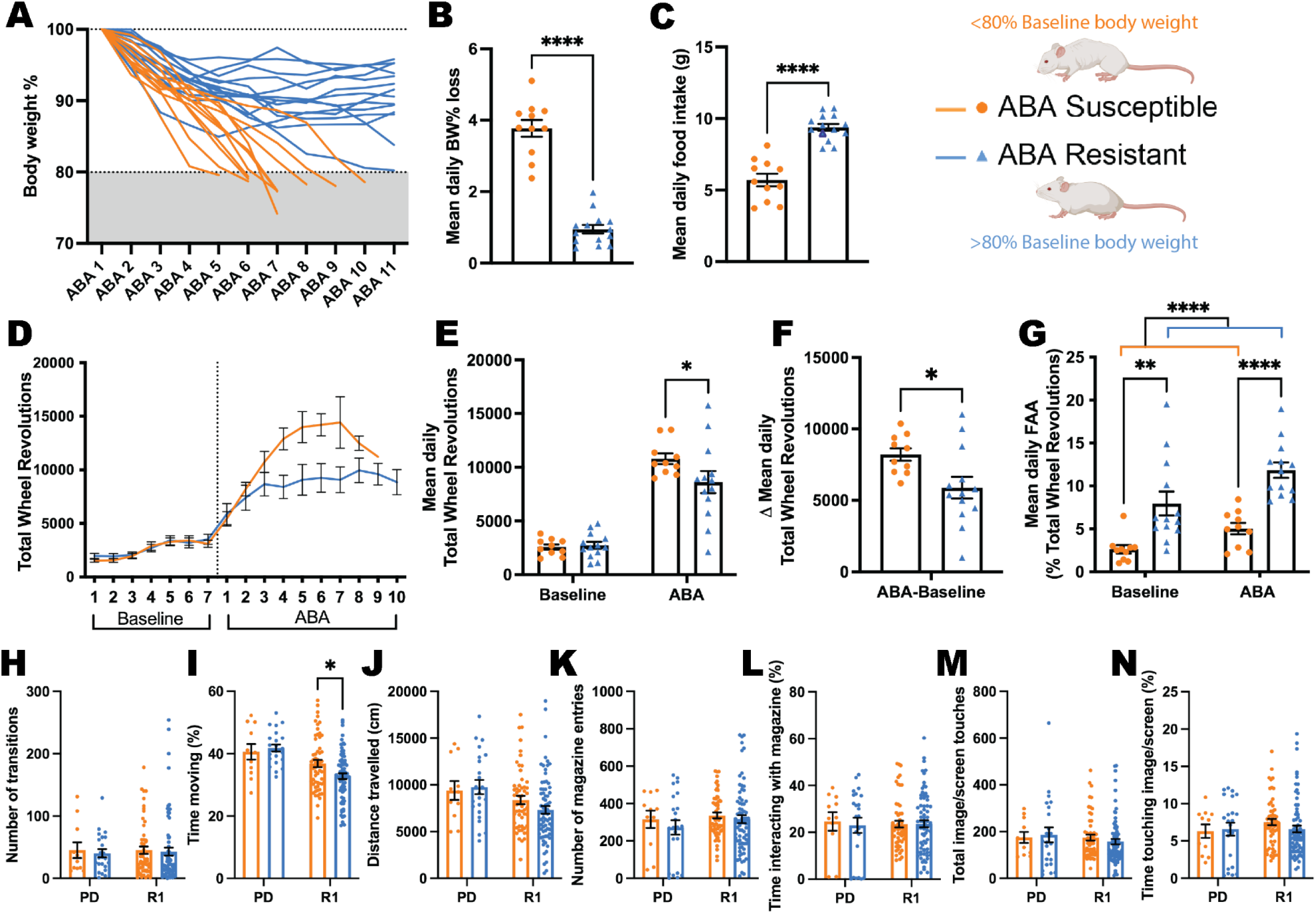
Key activity-based anorexia (ABA) parameters that differentiate individuals that are susceptible and resistant to ABA and behavioural profiles during cognitive testing. Data are from animals that underwent ABA <u>after</u> cognitive testing in the PhenoSys. **(A)**Body weight (% of baseline) trajectories for individual animals showing that animals split into two subpopulations: ABA susceptible or ABA resistant. **(B)** Mean daily ABA body weight (BW) % loss (*p*<.0001). **(C)** Mean daily ABA food intake (*p*<.0001). **(D)** Daily running wheel activity (RWA) across both experimental phases. **(E)** Mean daily RWA (outcome*phase interaction *p*=.0232). ABA phase: ABA susceptible > ABA resistant (*p*=.0497). **(F)** Change in mean daily RWA from baseline to ABA (*p*=.0232). **(G)** Mean daily food anticipatory activity (FAA; RWA in the hour before food access; outcome *p*<.0001). ABA resistant > ABA susceptible during both baseline (*p*=.0011) and ABA (*p*<.0001). **(H)** Number of transitions into the PhenoSys touchscreen testing chamber. **(I)** Percentage of time moving in the chamber. R1: ABA susceptible > ABA resistant (*p*=.0101). **(J)** Distance (cm) travelled in the chamber. **(K)** Number of pellet magazine entries. **(L)** Percentage of time interacting with pellet magazine. **(M)** Number of total image/screen touches. **(N)** Percentage of time touching the image/screen. Bar graphs show group mean ± SEM with individual animals (symbols); line graph shows group mean ± SEM. \**p*<.05, \*\**p*<.01, \*\*\*\**p*<.0001. PD: pairwise discrimination; R1: reversal learning. For full statistical analysis details and results see **Supplementary Table 2**.

**Supplementary Figure 8.**
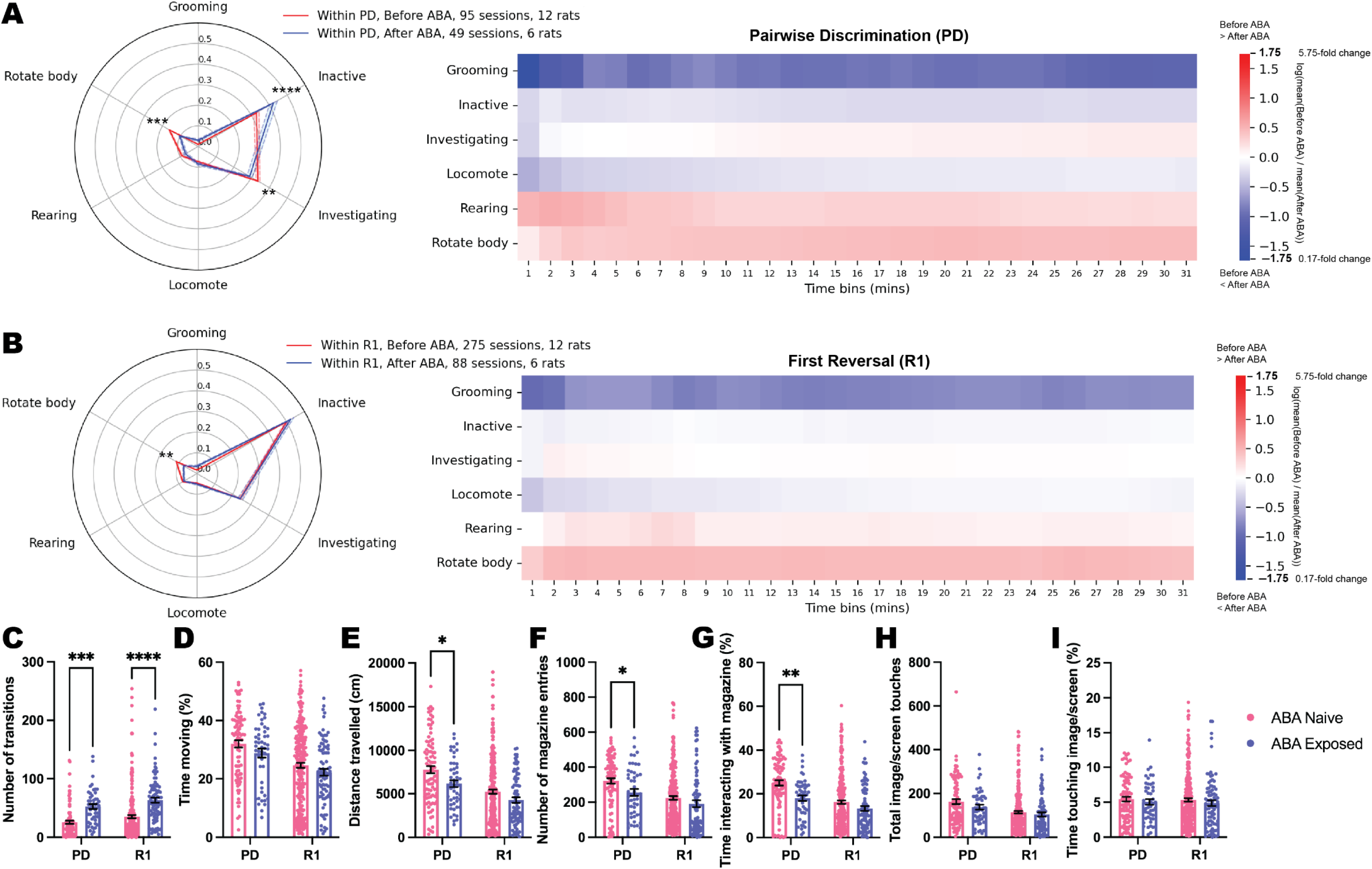
Behavioural differences during touchscreen testing due to order effects of PhenoSys and activity-based anorexia (ABA) exposure. Spider plots and heat maps show the proportion of time spent doing each behaviour within each session video during pairwise discrimination (PD; **A**) or first reversal (R1; **B**). The spider plots show group mean ± SEM (shaded bands). The time bin heat maps show the change in these proportion values between the groups across time. The values are the log(mean(Before ABA)/mean(After ABA)), where log is the natural log. The time bins are cumulative, showing e.g. 0-1 mins, 0-2 mins, etc. **(A)** Within PD (behaviour*ABA timing interaction *p*<.0001), the After ABA rats spent significantly more time inactive (*p*<.0001), and significantly less time rotating their body (*p*=.0002) and investigating (*p*=.0078) than the Before ABA group. **(B)** Within R1 (behaviour*ABA timing interaction *p*=.0028), the After ABA rats spent significantly less time rotating their body than the Before ABA rats (*p*=.0039). **(C-I)** Bar graphs show group mean ± SEM with individual sessions (symbols); ABA Naive = PhenoSys Before ABA; ABA Exposed = PhenoSys After ABA. **(C)** Number of transitions in the chamber (ABA exposure *p*<.0001). ABA Exposed > ABA Naive during both PD (*p*=.0001) and R1 (*p*<.0001). **(D)** Percentage of time moving in the chamber. **(E)** Distance (cm) travelled in the chamber (ABA exposure *p*=.0014). PD: ABA Naive > ABA Exposed (*p*=.0277). **(F)** Number of pellet magazine entries (ABA exposure *p*=.0053). PD: ABA Naive > ABA Exposed (*p*=.0454). **(G** Percentage of time interacting with pellet magazine (ABA exposure *p*<.0001). PD: ABA Naive > ABA Exposed (*p*=.0014). **(H)** Number of total image/screen touches. **(I)** Percentage of time touching the image/screen. \**p*<.05, \*\**p*<.01, \*\*\**p*<.001, \*\*\*\**p*<.0001. For full statistical analysis details and results see **Supplementary Table 2**.

**Supplementary Figure 9.**
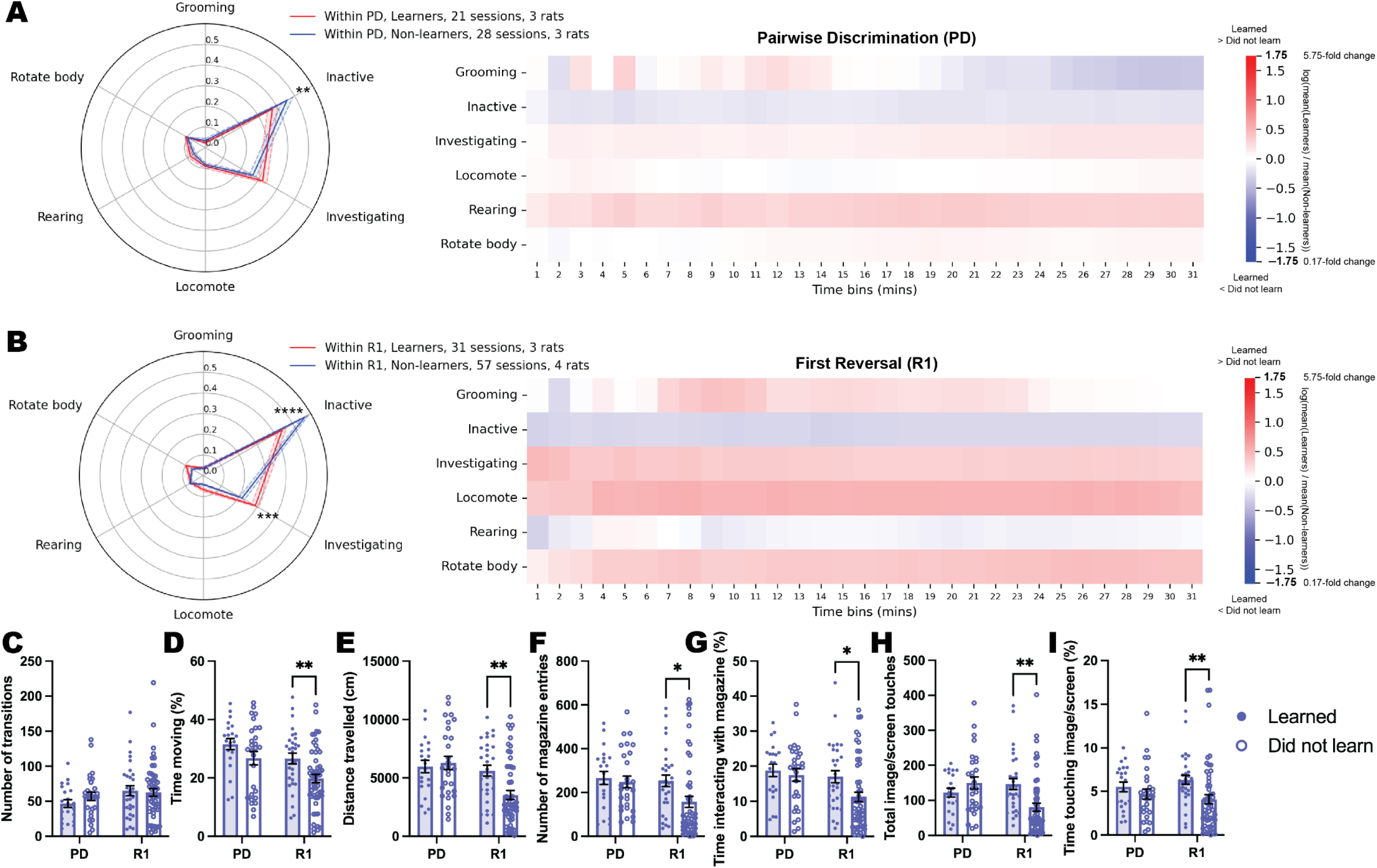
Behavioural differences during touchscreen testing due to whether rats learned or did not learn first reversal after prior exposure to ABA. Spider plots and heat maps show the proportion of time spent doing each behaviour within each session video during pairwise discrimination (PD; **A**) or first reversal (R1; **B**). The spider plots show group mean ± SEM (shaded bands). The time bin heat maps show the change in these proportion values between the groups across time. The values are the log(mean(Learners)/mean(Non-learners)), where log is the natural log. The time bins are cumulative, showing e.g. 0-1 mins, 0-2 mins, etc. **(A)** Within PD (behaviour*outcome interaction *p*=.0045), the non-learners spent significantly more time inactive than the learners (*p*<.0059). **(B)** Within R1 (behaviour*outcome interaction *p*<.0001), the non-learners spent significantly more time inactive (*p*<.0001) and significantly less time investigating (*p*=.0006) than the learners. **(C-I)** Bar graphs show group mean ± SEM with individual sessions (symbols). **(C)** Number of transitions in the chamber. **(D)** Percentage of time moving in the chamber (outcome *p*=.0031). R1: ABA Exposed learned > did not learn (*p*=.0100). **(E)** Distance (cm) travelled in the chamber (stage*outcome interaction *p*=.0185). R1: ABA Exposed learned > did not learn (*p*=.0021). **(F)** Number of pellet magazine entries (outcome *p*=.0557). R1: ABA Exposed learned > did not learn (*p*=.0199). **(G)** Percentage of time interacting with pellet magazine (outcome *p*=.0448). R1: ABA Exposed learned > did not learn (*p*=.0172). **(H)** Number of total image/screen touches (stage*outcome interaction *p*=.0025). R1: ABA Exposed learned > did not learn (*p*=.0012). **(I)** Percentage of time touching the image/screen (outcome *p*=.0136). R1: ABA Exposed learned > did not learn (*p*=.0075). \**p*<.05, \*\**p*<.01, \*\*\**p*<.001, \*\*\*\**p*<.0001. For full statistical analysis details and results see **Supplementary Table 2**.

**Supplementary Figure 10.**
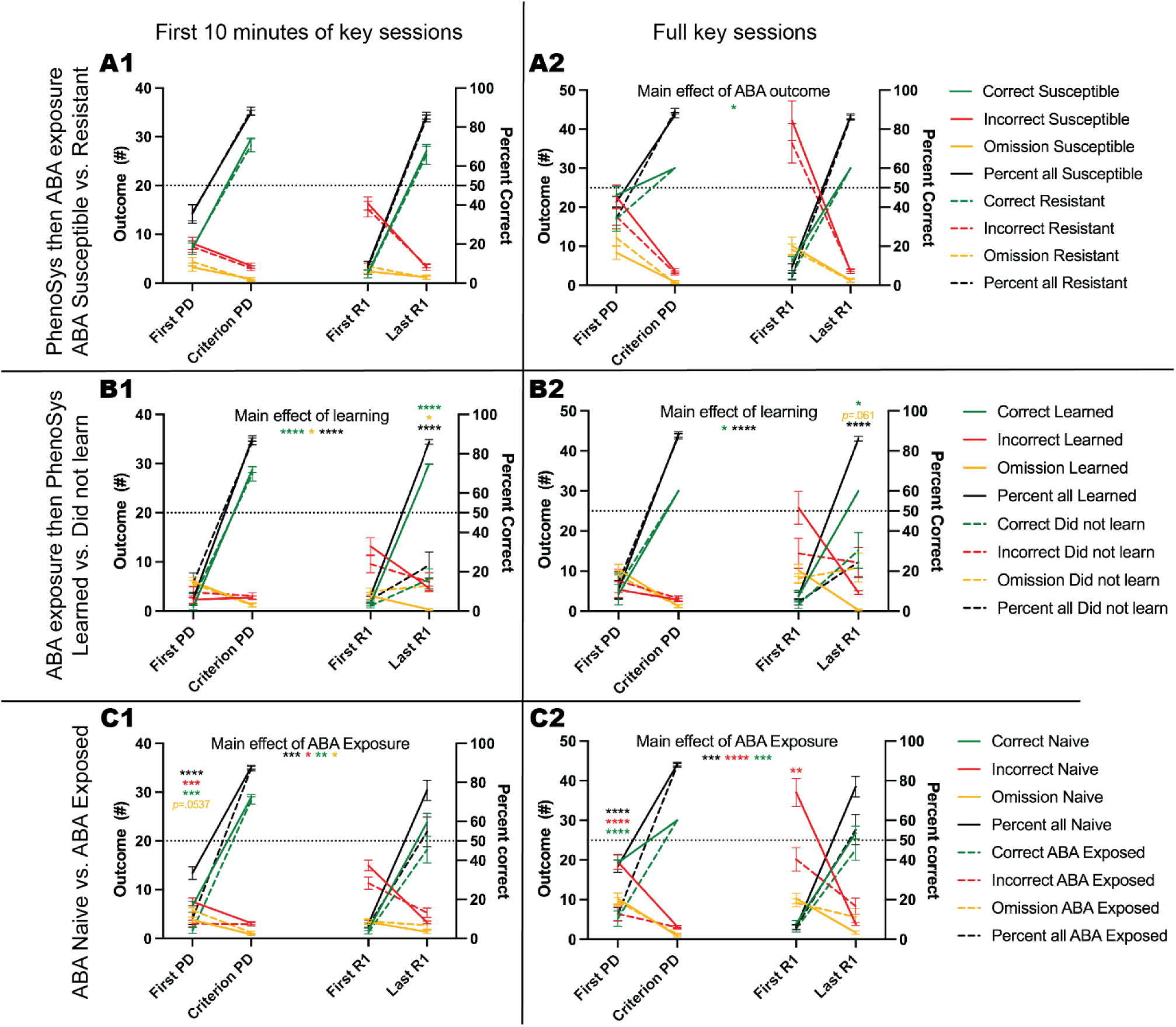
Performance during the first and last pairwise discrimination (PD) and first reversal (R1) sessions. Performance during the first 10 minutes **(1)** or the full **(2)** session for each critical session (see below) comparing between ABA Naive animals that were susceptible or resistant to ABA **(A)**, ABA Exposed animals that learned or did not learn R1 **(B)**, and ABA Naive versus ABA Exposed animals **(C)**. Critical sessions are: **First PD**, First session of PD, this is the animal’s first exposure to the two novel stimuli and the pairwise discrimination task; **Criterion PD,** The session of PD in which an animal reached progression criterion (i.e. Made 30 correct responses with >80% accuracy); **First R1,** The first session of R1, this is each animal’s first exposure to the reversed reward contingencies; and **Last R1,** The last session of R1, for animals that successfully learned R1 within 20 sessions this is the session in which they reached progression criterion (i.e. made 30 correct responses with <80% accuracy), for animals that did not reach R1 progression criterion this is their last session (i.e. session 20). **A2)** Correct trials, ABA outcome *p*=.0486. **B1)** <u>Correct trials</u>, all *ps*<.0001, Last R1 Learned > Did not learn *p*<.0001. <u>Omission trials</u>, all *ps*<.0192, Last R1 Learned < Did not learn *p*=.0120. <u>Percent correct</u>, all *ps*<.0001, Last R1 Learned > Did not learn *p*<.0001. **B2)** <u>Correct trials</u>, all *ps*<.0443, Last R1 Learned > Did not learn *p*=.0312. <u>Incorrect trials</u>, outcome*session interaction *p*=.0090. <u>Omission trials</u>, outcome*session interaction *p*=.0006, Last R1 Learned < Did not learn *p*=.0610. <u>Percent correct</u>, all *ps*<.0001, Last R1 Learned > Did not learn *p*<.0001. **C1)** <u>Correct trials</u>, ABA exposure *p*=.0024, First PD ABA Naive > ABA Exposed *p*=.0001. <u>Incorrect trials</u>, all *ps*<.0159, ABA Naive > ABA Exposed *p*=.0005. <u>Omission trials</u>, ABA exposure *p*=.0168, ABA Naive < ABA Exposed *p*=.0537. <u>Percent correct</u>, all *ps*<.0007, First PD ABA Naive > ABA Exposed *p*<.0001. **C2)** <u>Correct trials</u>, all *ps*<.0003, First PD ABA Naïve > Aba exposed *p*<.0001. <u>Incorrect trials</u>, all *ps*<.0001, First PD ABA Naive > ABA Exposed *p*<.0001, First R1 ABA Naive > ABA Exposed *p*=.0024. <u>Percent correct</u>, all *ps*<.0002, First PD ABA Naive > ABA Exposed *p*<.0001. Data are group mean ± SEM. \**p*<.05, \*\**p*<.01, \*\*\**p*<.001, \*\*\*\**p*<.0001. For full statistical analysis details and results see **Supplementary Table 2**.

**Supplementary Table 1.** Parameters for pretraining and 2VDLR in the novel touchscreen apparatus

| Stages | Response | Trial Outcome | Reward | Trial Initiation | Progression Criterion |
| --- | --- | --- | --- | --- | --- |
| **Habituation** | N/A | N/A | Sucrose pellets spread throughout PhenoSys | N/A | Consumption of 15-20 pellets in testing chamber |
| **Initial Touch** | Touch image | Correct tone + magazine LED illumination | 2 sucrose pellets | Automatic start after 10s ITI end | 2 sessions with 30 (maximum) trials |
|  | Touch to blank window during image presentation |  | 1 sucrose pellet |  |  |
|  | No screen touch |  |  |  |  |
| **Must Touch** | Touch image | Correct tone + magazine LED illumination | 1 sucrose pellets | Automatic start after 10s ITI end | 2 sessions with 30 (maximum) trials |
|  | Touch to blank window during image presentation |  |  |  |  |
| **Must Initiate** | Touch image | Correct tone + magazine | 1 sucrose pellets | Must be initiated by magazine nose | 2 sessions with 30 (maximum) trials |
|  | Touch to blank window during image presentation |  |  |  |  |
|  |  | LED illumination |  | poke after 10s<br>ITI end |  |
| Punish Incorrect | Touch image | Correct tone +<br>magazine LED illumination | 1 sucrose pellets | Must be initiated by<br>magazine nose poke after 10s<br>ITI end | 2 sessions with 30 (maximum) trials with ≥80% accuracy |
|  | Touch to blank window during image presentation | Incorrect tone +<br>house light on + 5s time out | No reward | Must be initiated by<br>magazine nose poke after 5s time out + 15s<br>ITI end |  |
|  | No screen touch |  |  |  |  |
| Pairwise Discrimination +<br>Reversal Learning | Touch to correct image | Correct tone +<br>magazine LED illumination | 1 sucrose pellets | Must be initiated by<br>magazine nose poke after 10s time out + 10s<br>ITI end | 2 sessions with 30 (maximum) trials with ≥80% accuracy |
|  | Touch to incorrect image | Incorrect tone +<br>house light on + 10s time out | No reward |  |  |
|  | No screen touch |  |  |  |  |

**ITI:** inter-trial interval.

**Supplementary Table 2.**
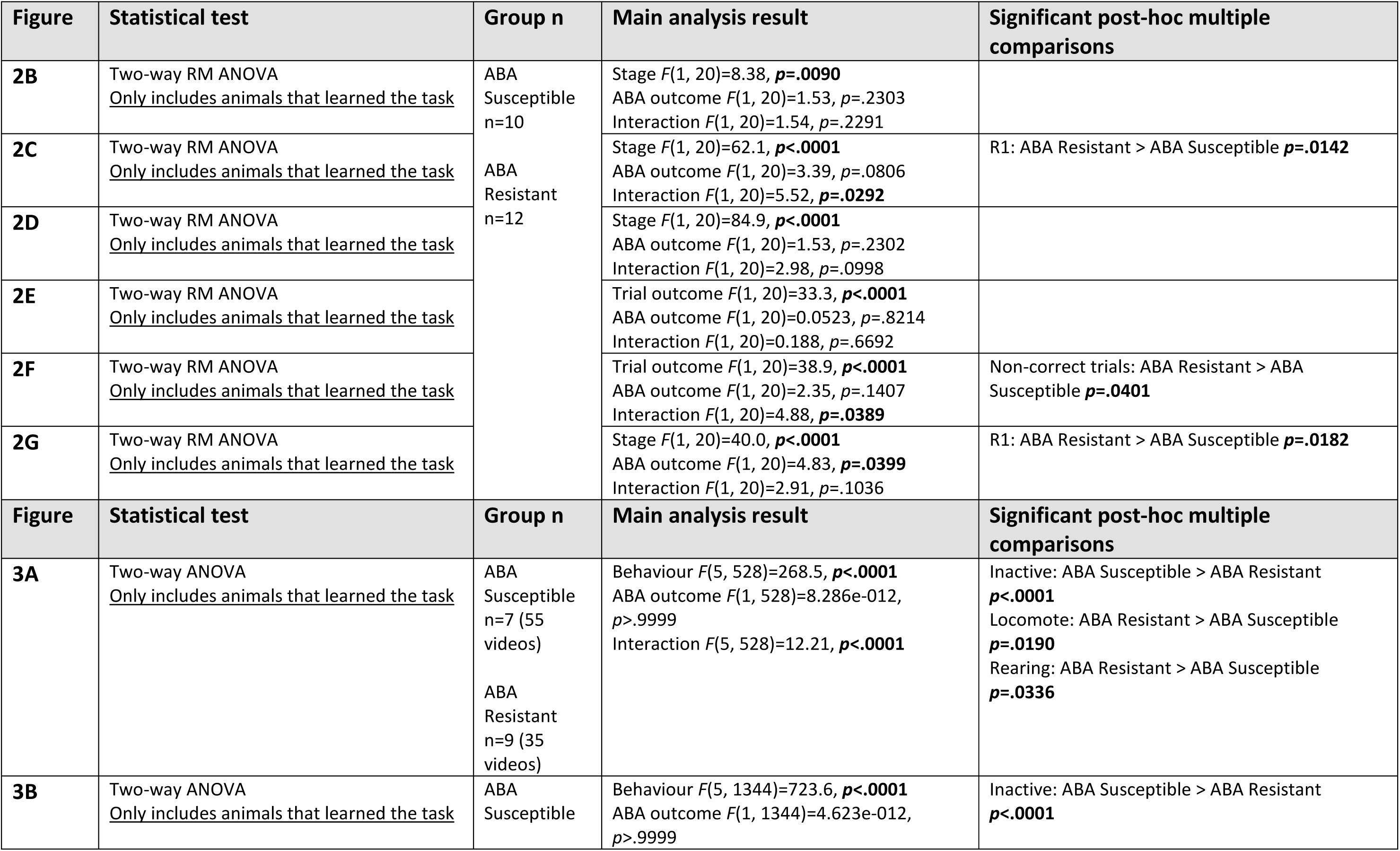
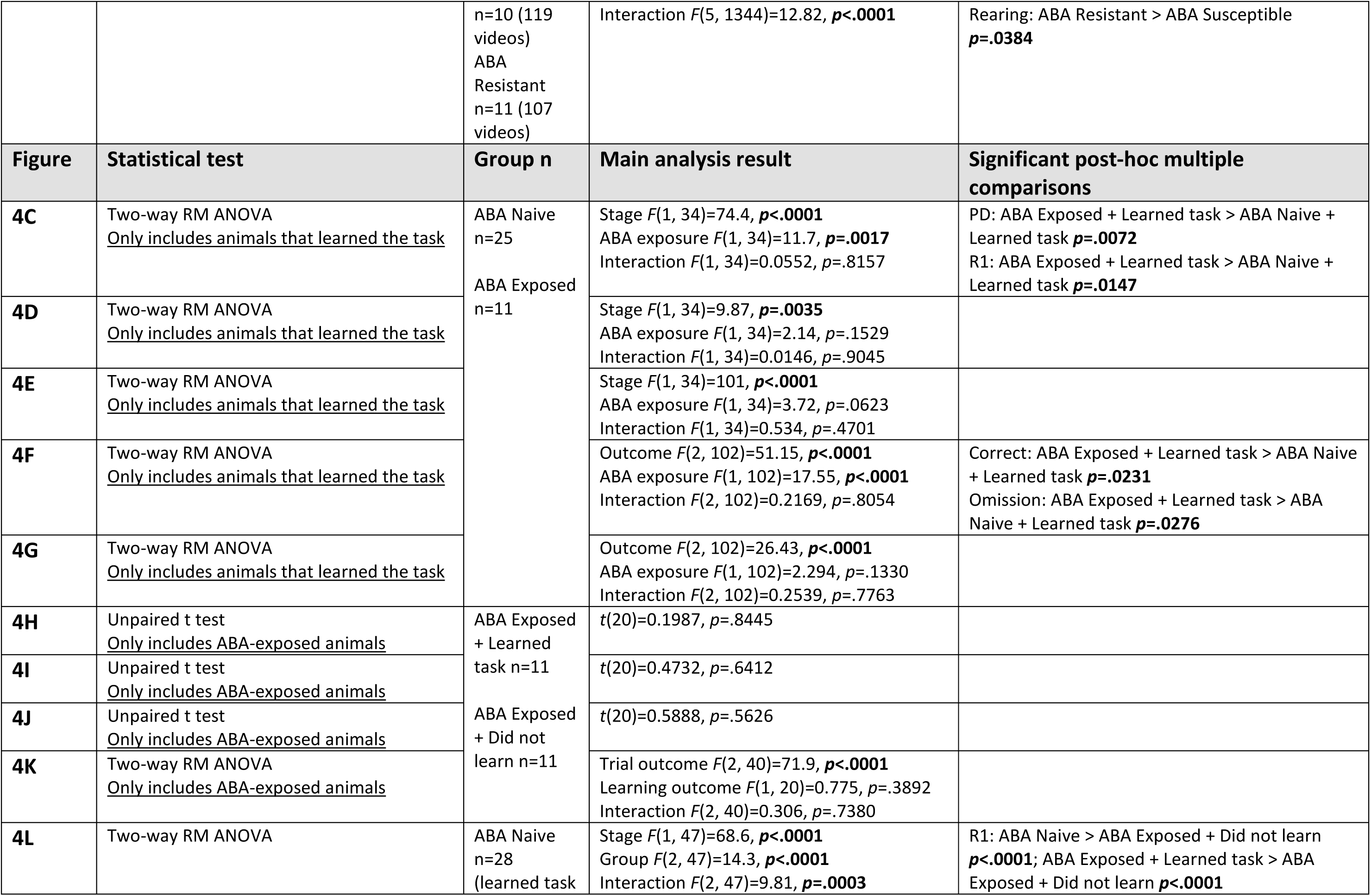
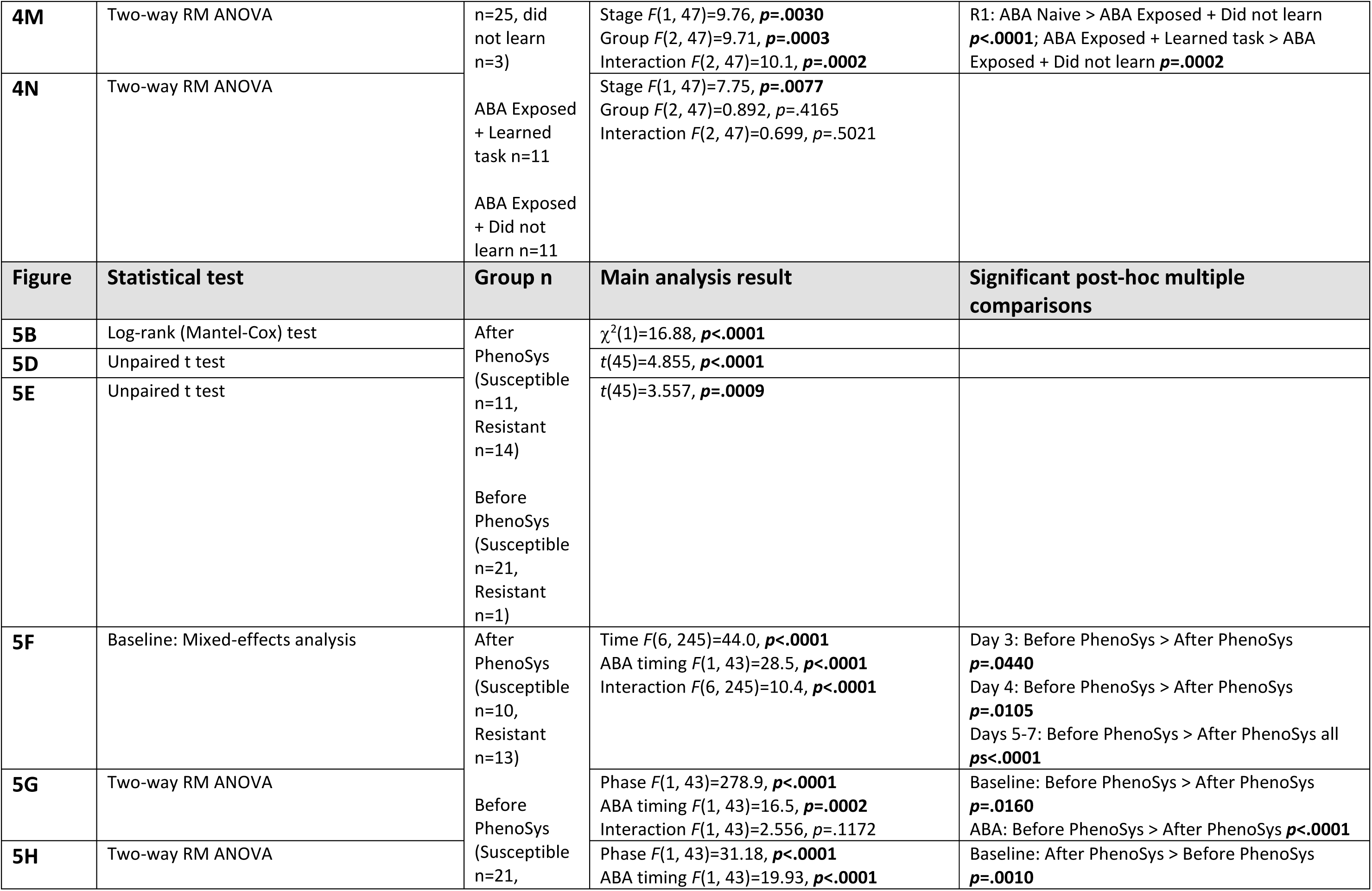
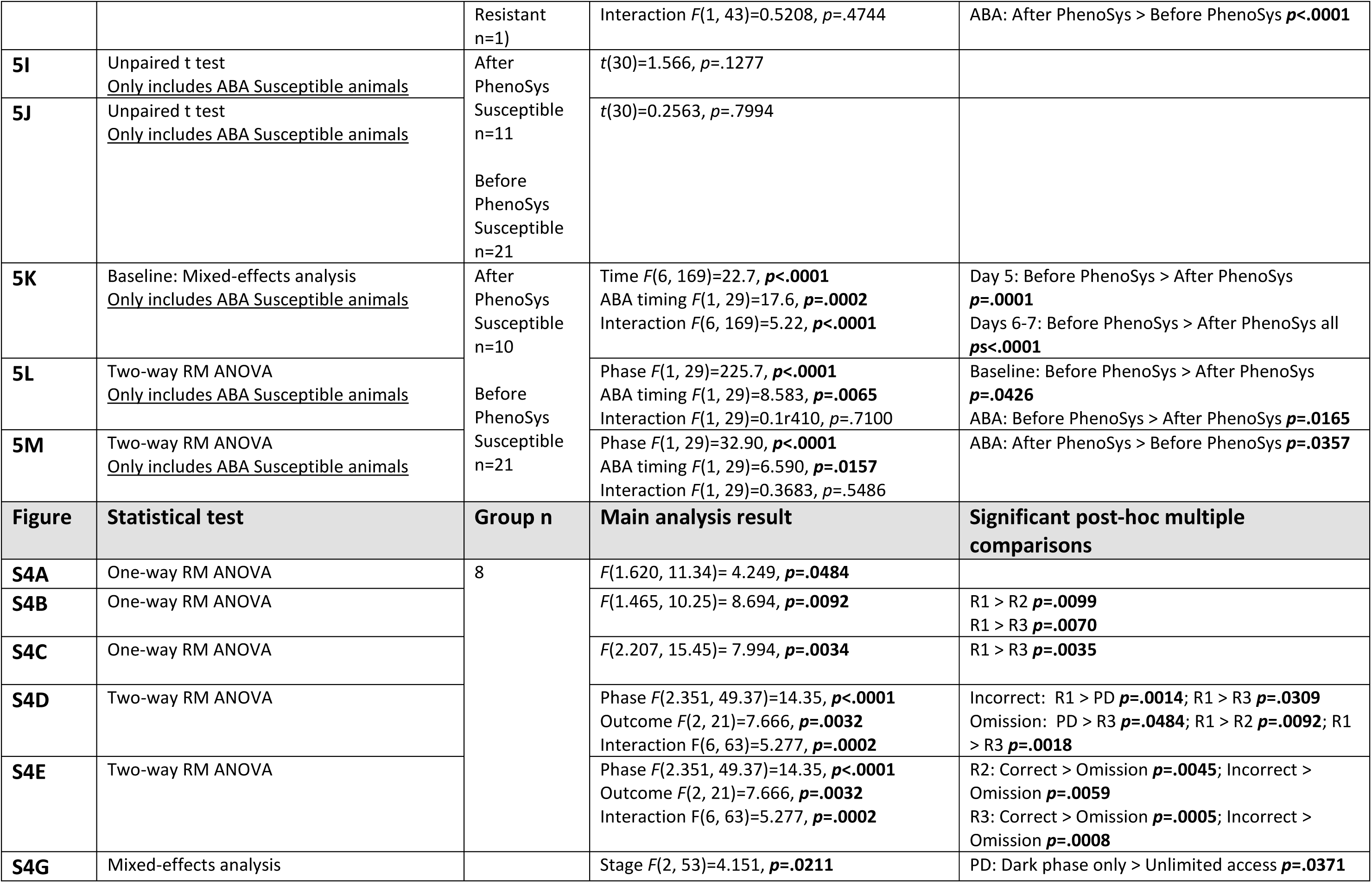
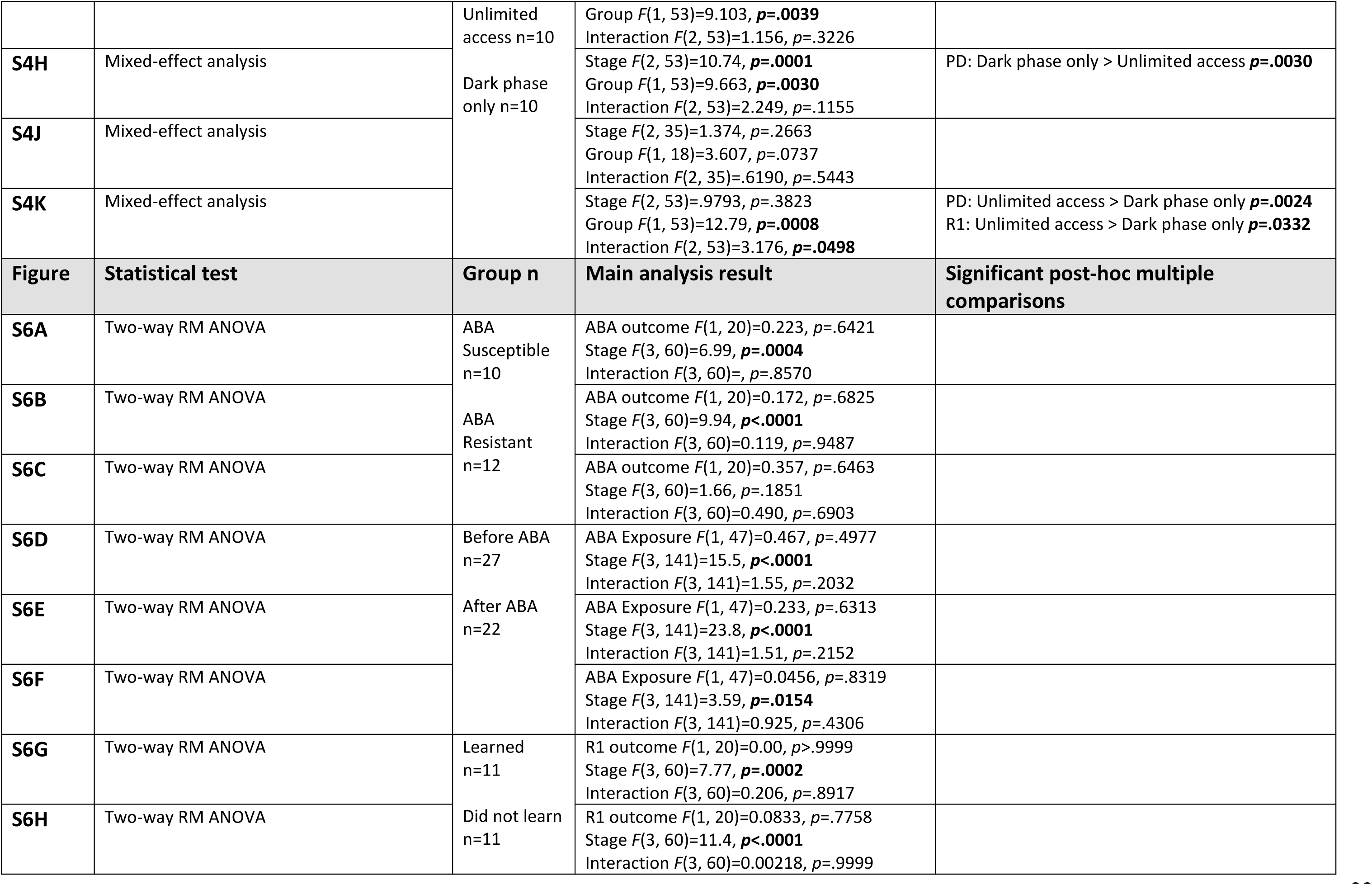
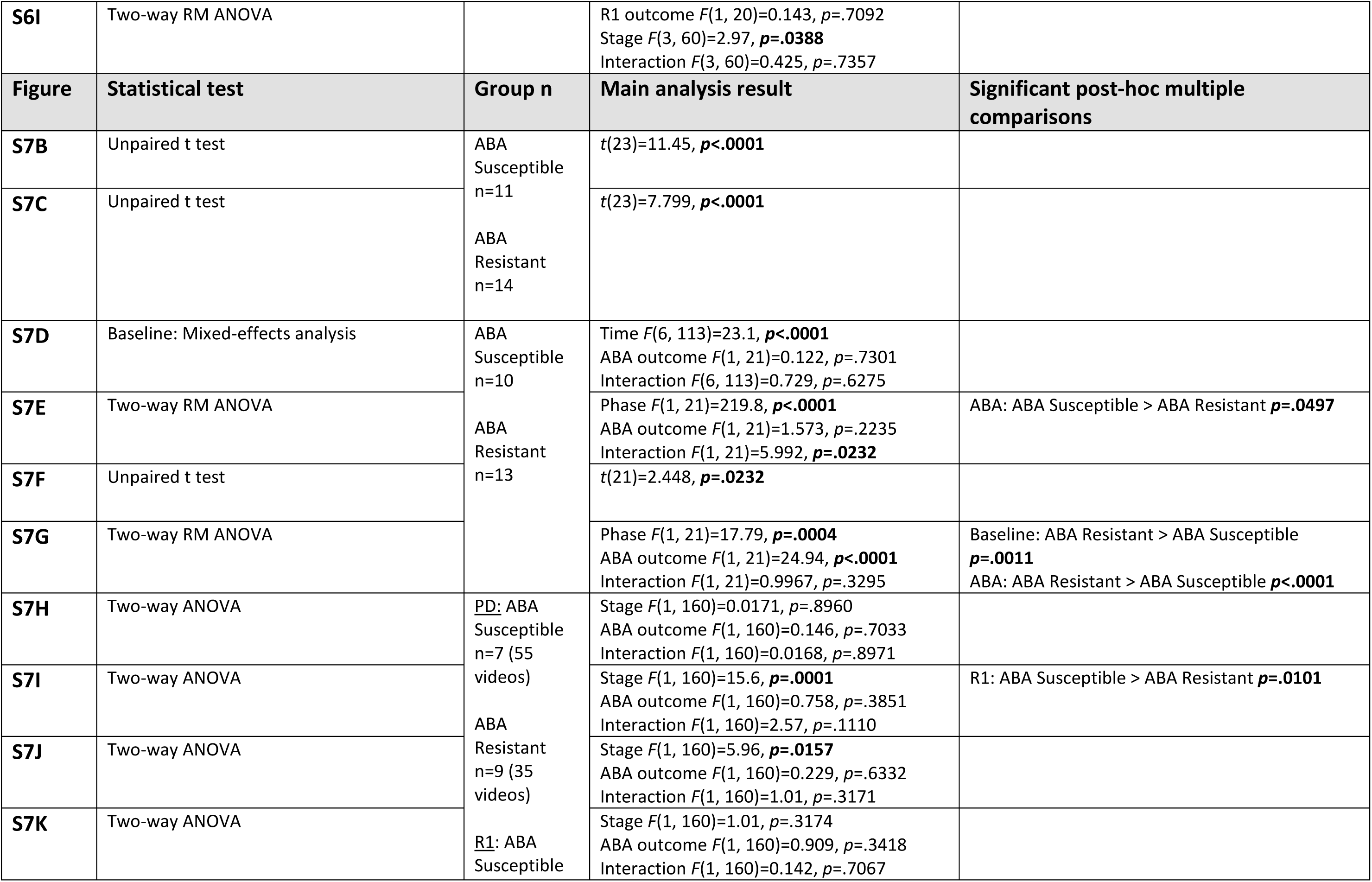
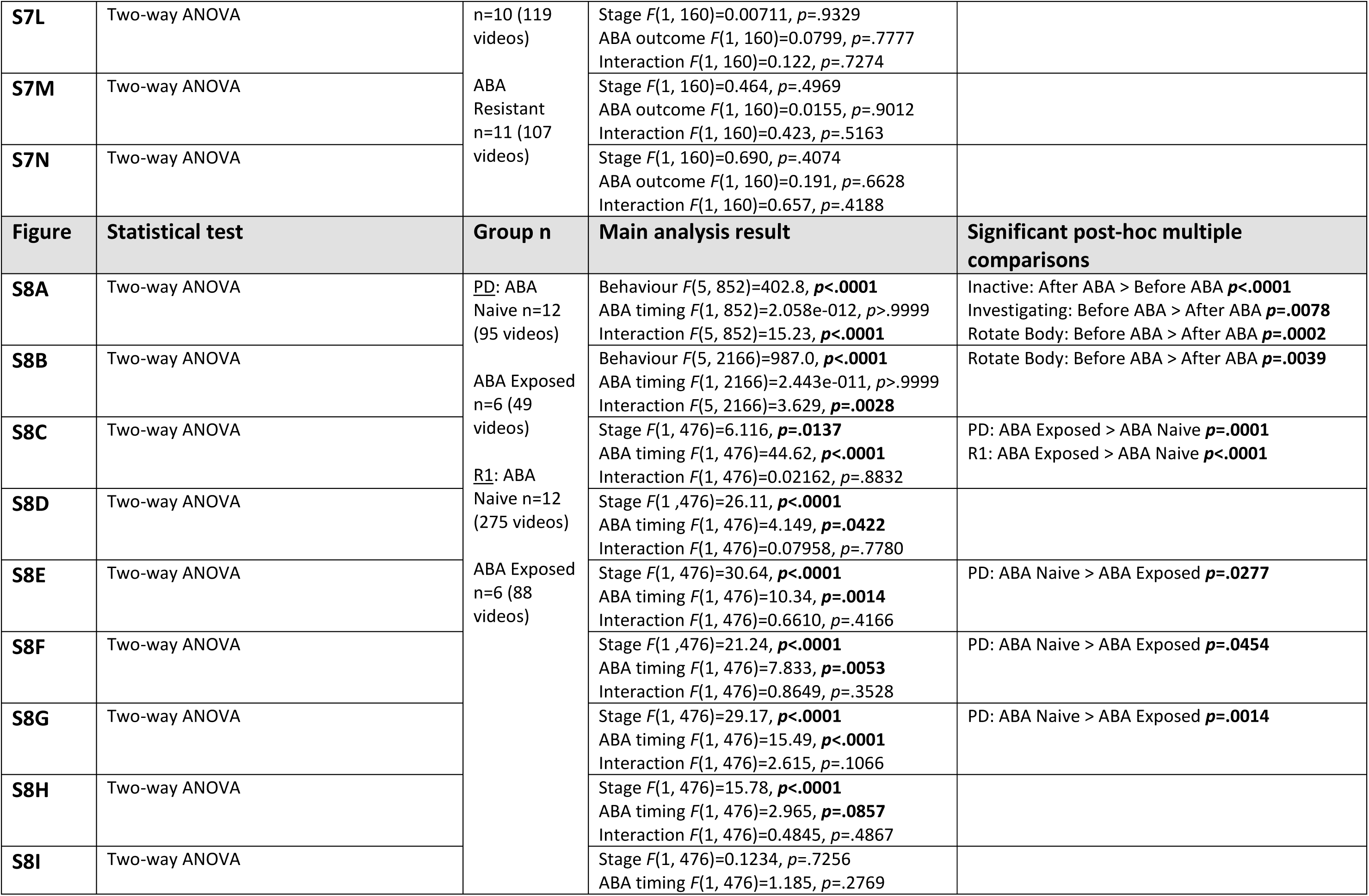
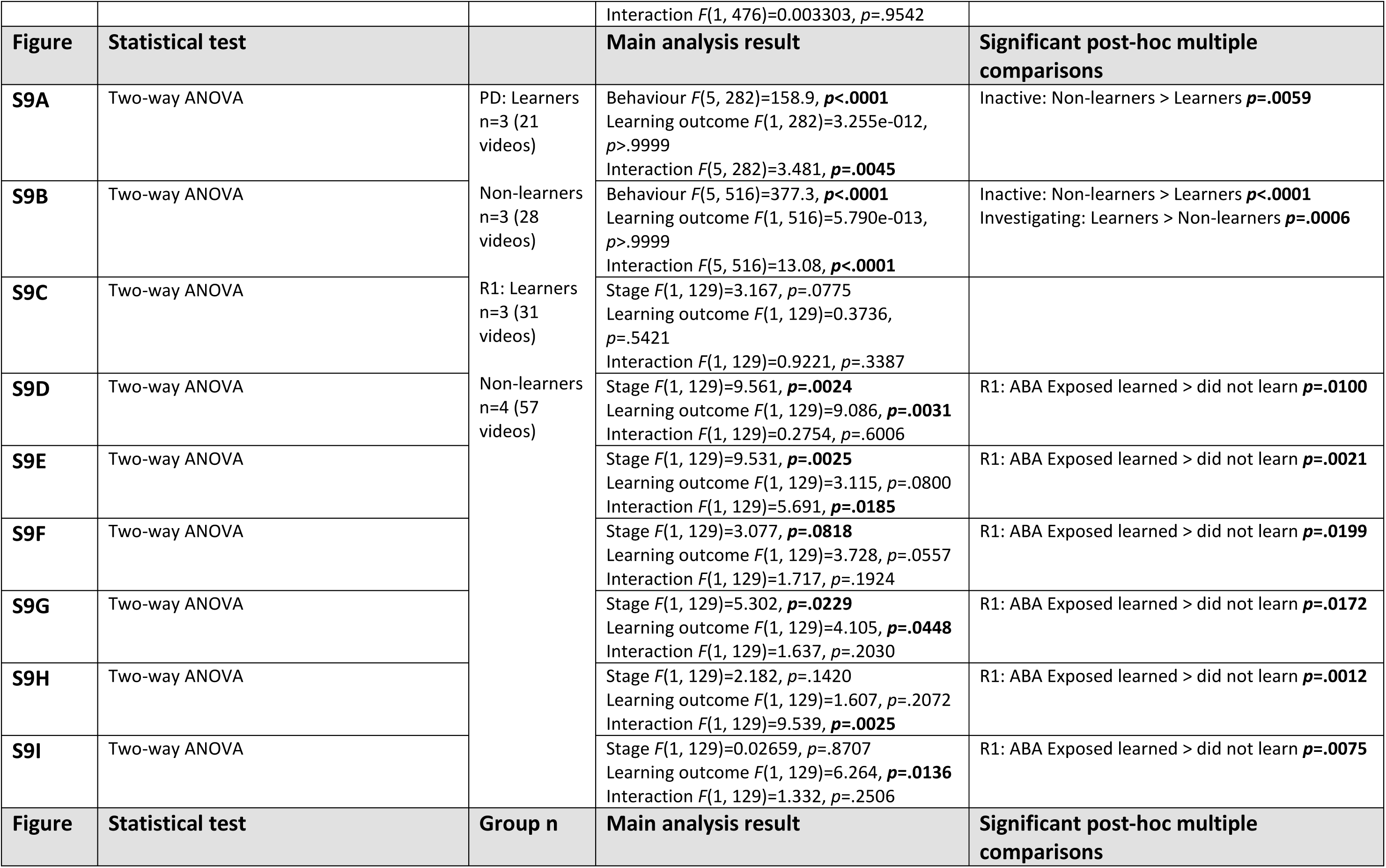
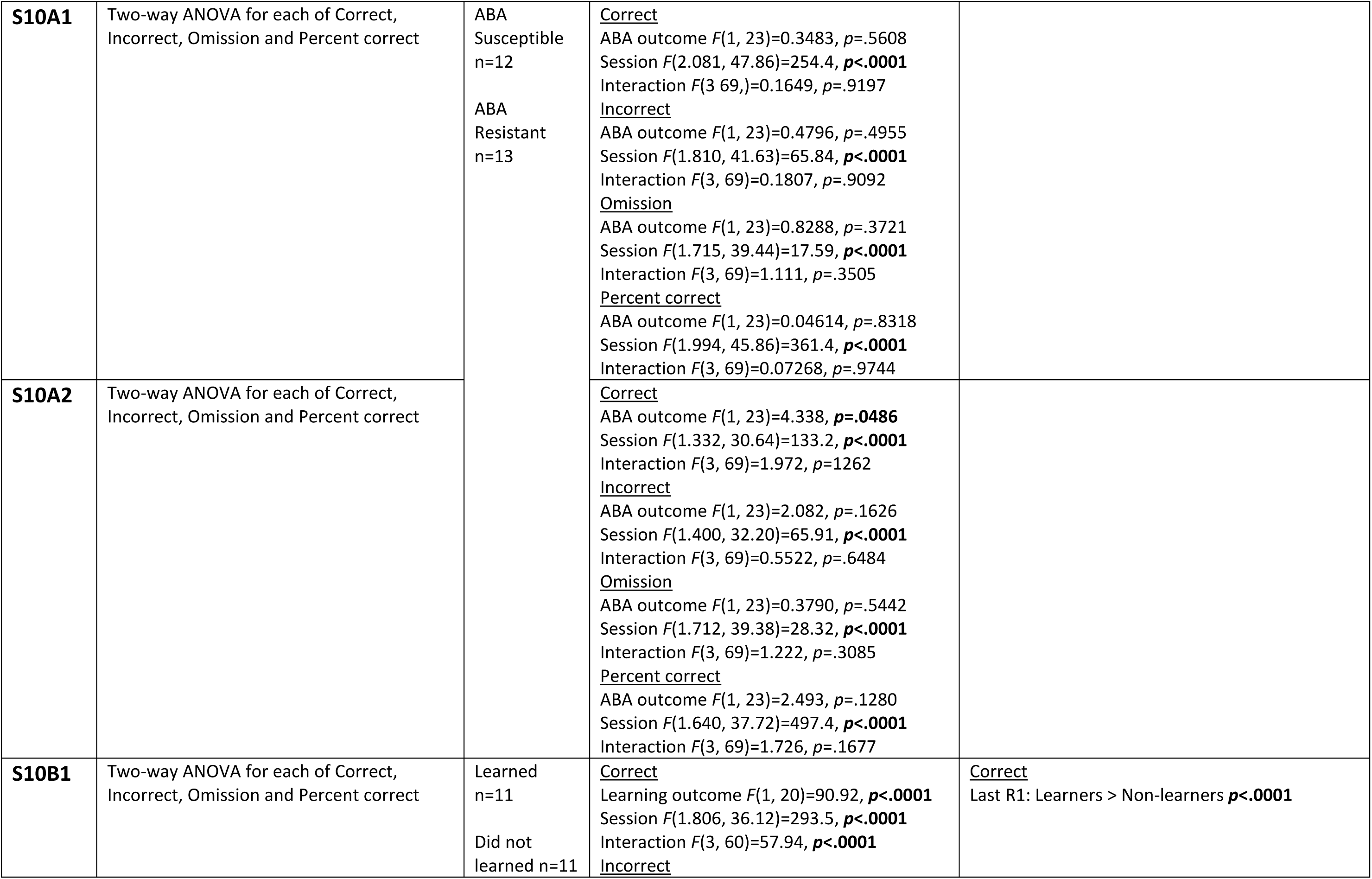
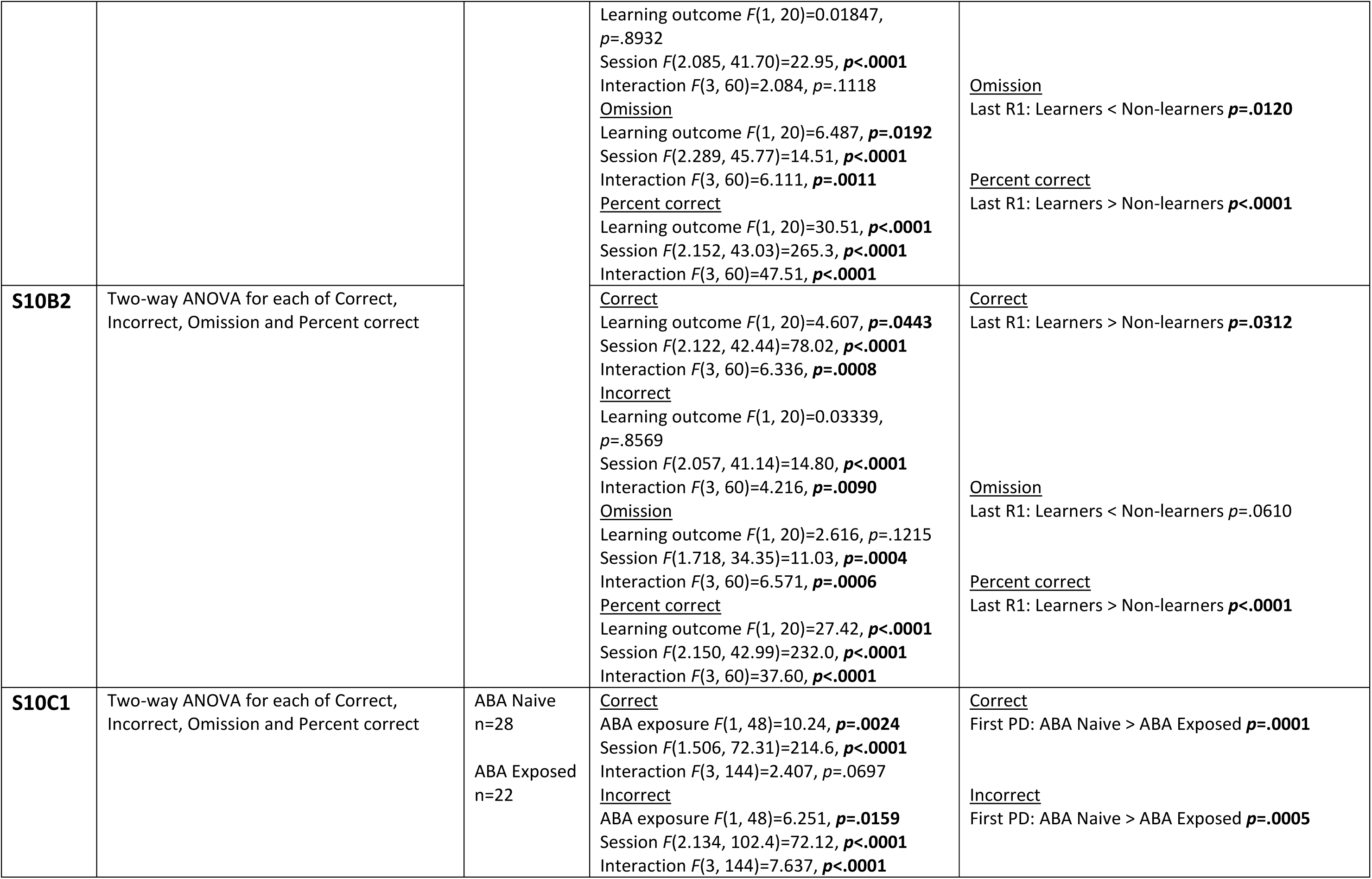
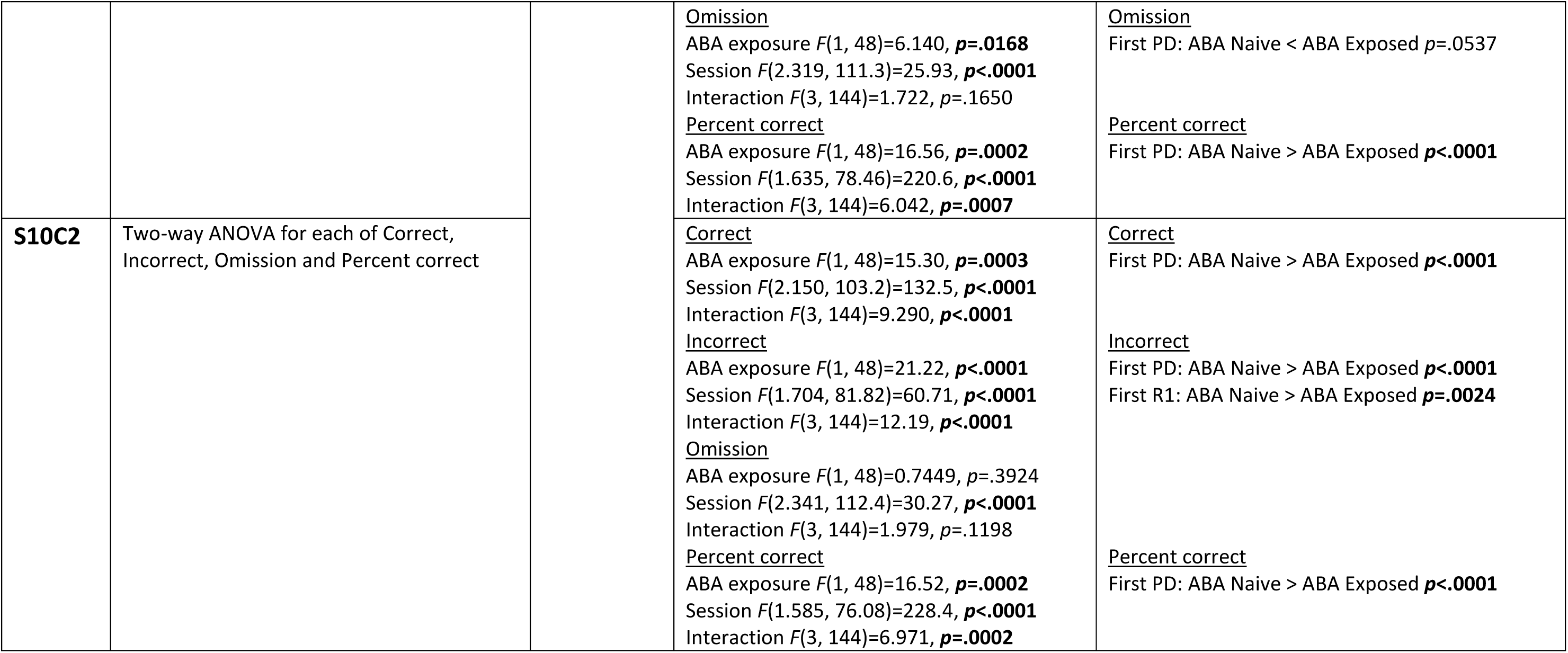
Statistical test details and results for all analyses

## Notes

https://github.com/Foldi-Lab/PhenoSys-data

https://github.com/Foldi-Lab/PhenoSys-codes

https://doi.org/10.6084/m9.figshare.21539685

### Competing Interest Statement

The authors have declared no competing interest.

